# Resolving Hepatic Metabolic Flux During Ex Vivo Liver Perfusion Reveals Compartment-Specific Constraints

**DOI:** 10.64898/2026.11.24.769061

**Authors:** Amin M. Amin, Marc McLeod, Nicholas Browder, Melissa Inigo-Vollmer Massey, Justin Fletcher, Shawn C. Burgess

## Abstract

Normothermic machine perfusion (NMP) is used to preserve and assess donor livers, but current viability criteria rely on global measures that do not resolve intracellular metabolism, which would provide a more mechanistic measure of donor liver function. We combined staged ex vivo perfusion of mouse livers, serial lobectomy and ¹³C tracers ([U-¹³C₃]lactate, [U-¹³C₃]pyruvate and [U-¹³C₆]glucose) to test whether glucose fragment-based flux models used in vivo describe intact hepatic metabolism. With intracellular metabolite isotopomers as constraints, a homogeneous one-compartment model failed to fit most datasets, and the lack of fit centered on TCA cycle intermediates such as malate. A two-compartment model, in which a glycolytic compartment takes up glucose from the sinusoidal pool, including glucose released by a gluconeogenic compartment, fitted 41 of 45 datasets. Flux partitioning between compartments depended on nutritional state, which influenced hepatic metabolism more than perfusate glucose. Nevertheless, glucose fragment and two-compartment flux estimates correlated (R² = 0.68–0.89). Intracellular isotope tracing thus shows that hepatic compartmentation constrains whole-liver flux estimation, while supporting simplified models for quantitative assessment during perfusion.

**HIGHLIGHTS:**

- Intracellular ¹³C labeling shows the perfused liver is not one metabolic compartment
- Data fit a gluconeogenic compartment that supplies glucose to a glycolytic one
- Donor nutritional state, not perfusate glucose, governs liver metabolism ex vivo
- Simple glucose fragment models still capture the dominant hepatic fluxes

## INTRODUCTION

Normothermic machine perfusion (NMP) is changing donor liver preservation from static cold storage into a platform for organ assessment, metabolic rehabilitation and therapeutic intervention before transplantation^1–3^. Viability is currently judged from integrated outputs such as lactate clearance, bile production, perfusion hemodynamics and markers of hepatocellular injury^2,4,5^. These measures show whether the liver is metabolically active, but not how its intracellular pathways are organized. Because coordinated oxidative metabolism, anaplerosis and gluconeogenesis are central to hepatic function^6,7^, direct measurement of these pathways could provide a more mechanistic assessment of the perfused liver.

Stable isotope tracing combined with metabolic flux analysis (MFA) is the reference approach for quantifying hepatic gluconeogenesis, anaplerosis, pyruvate cycling and tricarboxylic acid (TCA) cycle flux in vivo^8–12^. Because intracellular metabolites are rarely accessible in vivo, these approaches infer intrahepatic fluxes mainly from the positional and mass isotopomers of circulating glucose. GC–MS analysis of several derivatized glucose fragments, each reporting a defined subset of glucose carbons, now allows these measurements to be made from small plasma volumes^10,11,13^. Glucose fragment-based models built on these measurements have been used to quantify hepatic flux in a range of physiological and genetic models^12,14^.

To make flux estimation tractable from glucose labeling alone, these models represent the liver as a single, well-mixed compartment and simplify exchange reactions, pyruvate cycling and CO₂ recycling^11,15^. The liver is not homogeneous, however. Periportal and pericentral hepatocytes differ in their oxidative, gluconeogenic, glycolytic and nitrogen-handling programs^16–23^. Labeling can also equilibrate incompletely between cytosolic and mitochondrial metabolite pools^24–26^, and non-parenchymal cells contribute to tissue metabolite pools^27^. Isotopomer studies in perfused rat liver identified heterogeneous labeling of TCA cycle intermediates decades ago^25,26,28^. Whether this heterogeneity limits the models now used to estimate hepatic flux in vivo has not been tested directly.

Ex vivo liver perfusion makes this test possible. Intracellular metabolites can be sampled alongside effluent glucose while the architecture of the intact organ is preserved^25,29–31^. Measuring intracellular and extracellular labeling in the same liver therefore allows assumptions that cannot be evaluated in vivo to be tested directly.

Here, we combined staged ex vivo perfusion of mouse livers with serial lobectomy, parallel [U-¹³C₃]lactate/[U-¹³C₃]pyruvate and [U-¹³C₆]glucose labeling of the same liver, GC–MS isotopomer analysis and MFA to ask whether a one-compartment glucose fragment-based model can account for intracellular isotopomer measurements. It could not. The lack of fit was metabolite-specific, centering on TCA cycle intermediates such as malate, and was resolved by a two-compartment model comprising a gluconeogenic main compartment and a glycolytic secondary compartment, which fitted 41 of 45 datasets. The partitioning of flux between these compartments varied with nutritional state. We therefore interpret the failure of the one-compartment model not as a computational limitation but as evidence that intact hepatic metabolism is functionally compartmentalized. Nevertheless, a simplified glucose fragment model recovered the dominant flux relationships estimated by the two-compartment model. These findings show that hepatic metabolic compartmentation, long recognized from zonation and heterogeneous isotopomer labeling, is large enough to constrain whole-liver flux estimation, yet does not compromise the dominant fluxes recovered by simplified glucose fragment models. This supports isotope-based assessment of liver metabolic function during machine perfusion.

## RESULTS

### Staged ex vivo perfusion preserves nutritional-state-dependent hepatic oxygen consumption

To obtain intracellular and extracellular isotopomer measurements from the same organ, we established a single-pass perfusion system with three sequential 30-min stages (Figures 1A–1C). In Stage 1, livers were perfused without glucose in the presence of [U-¹³C₃]lactate (1.5 mM) and [U-¹³C₃]pyruvate (0.15 mM). In Stage 2, unlabeled glucose (5 or 20 mM) was added. In Stage 3, the tracer was switched to [U-¹³C₆]glucose at the same glucose concentration. A lobe was removed and freeze-clamped at the end of each stage, so that intracellular labeling could be compared across conditions within the same liver (Figure 1B). Livers from fed and fasted mice were studied with or without insulin (Figure 1C).

**Figure 1.**
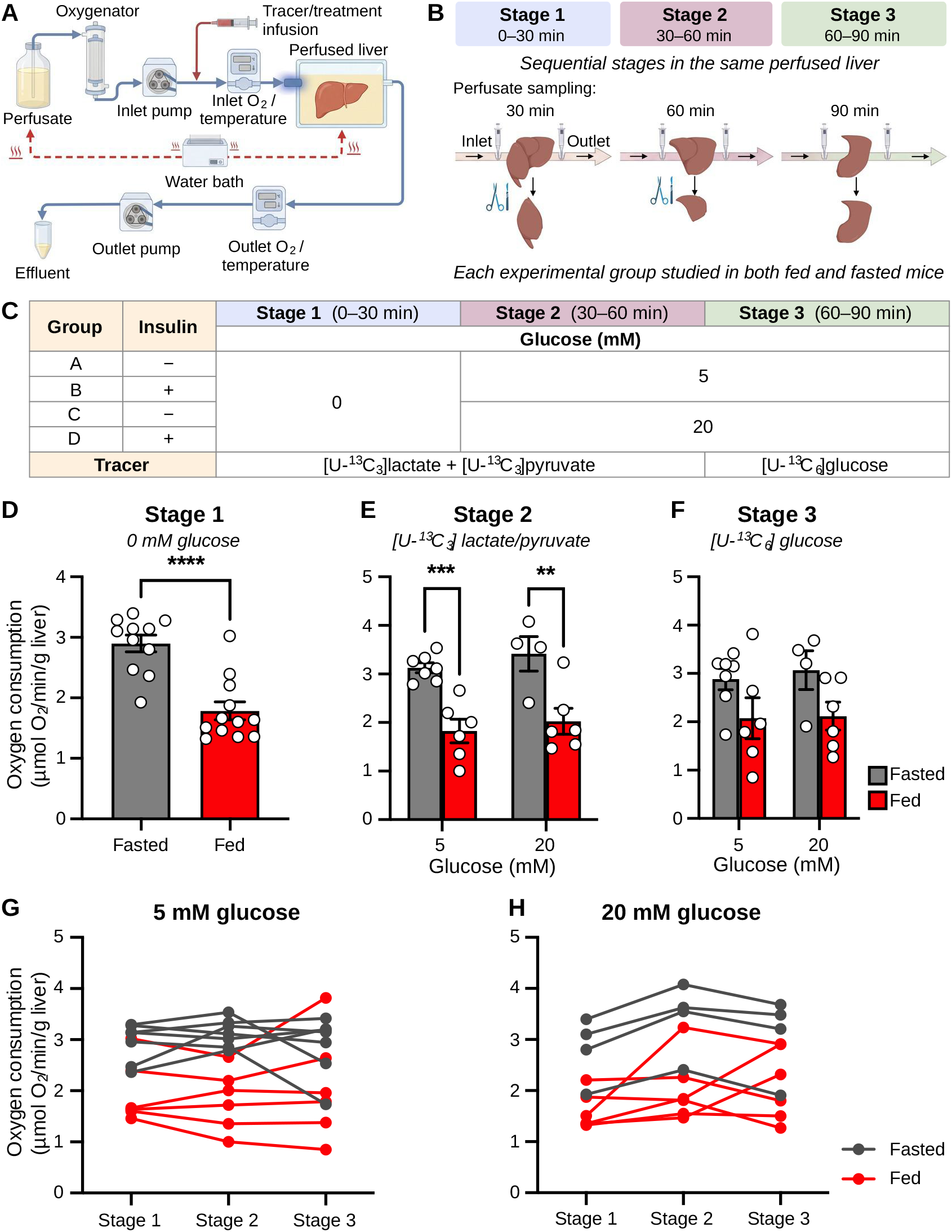
Stage-resolved ex vivo liver perfusion for isotope tracing shows that nutritional state, not perfusate glucose, determines hepatic oxygen consumption. (A) Single-pass (non-recirculating) perfusion circuit. Perfusate maintained at 37°C in a water-jacketed reservoir is pumped through a membrane oxygenator and inline oxygen and temperature sensors and enters the liver via the portal vein. Substrates and tracers are infused upstream of the liver by a programmable syringe driver. Oxygen tension is recorded continuously at the inlet and outlet to calculate hepatic oxygen consumption, and effluent is collected for metabolite and isotopomer analysis. (B) Perfusion protocol. Each liver was perfused for three consecutive 30-min stages. Effluent was sampled at 30, 60 and 90 min, and at the end of each stage a lobe was ligated, excised and snap-frozen in liquid nitrogen, allowing intracellular metabolites to be compared across stages within the same liver. (C) Experimental groups. Stage 1 was glucose-free with [U-^13^C_3_]lactate and [U-^13^C_3_]pyruvate as tracers. In Stage 2, unlabeled glucose (5 or 20 mM) was added with the same tracers. In Stage 3, the tracer was switched to [U-^13^C_6_]glucose at the same glucose concentration. Groups A–D differ by glucose concentration and the presence (+) or absence (−) of insulin; each group was studied in livers from fed and fasted mice. (D–F) Hepatic oxygen consumption in fasted (gray) and fed (red) livers during Stage 1 (D; 0 mM glucose), Stage 2 (E) and Stage 3 (F) at 5 or 20 mM glucose. (G and H) Oxygen consumption across stages in individual livers perfused at 5 mM (G) or 20 mM (H) glucose; lines connect measurements from the same liver. Data are mean ± SEM with individual livers shown. Insulin-treated and untreated livers were pooled because insulin had no significant effect (see Methods). n = 11 fasted and 12 fed livers in Stage 1 (D); n = 7 fasted and 6 fed livers at 5 mM and n = 4 fasted and 6 fed livers at 20 mM glucose (E–H). Oxygen consumption is expressed as µmol O_2_/min/g liver. (D) Welch’s unpaired t test. (E and F) Two-way ANOVA (nutritional state × glucose concentration) with uncorrected Fisher’s LSD test. Stage 2: nutritional state, *p* < 0.0001; glucose, *p* = 0.3118; interaction, *p* = 0.8603; fasted versus fed, *p* = 0.0005 (5 mM) and *p* = 0.0010 (20 mM). Stage 3: nutritional state, *p* = 0.0168; glucose, *p* = 0.7332; interaction, *p* = 0.8362. (G and H) Two-way repeated-measures ANOVA (nutritional state × perfusion stage) with Geisser–Greenhouse correction and Šídák’s multiple-comparisons test. Nutritional state, *p* = 0.0051 (5 mM) and *p* = 0.0091 (20 mM); stage, *p* = 0.9569 and *p* = 0.0613; interaction, *p* = 0.2536 and *p* = 0.5334. \**p* < 0.05, \*\**p* < 0.01, \*\*\**p* < 0.001, \*\*\*\**p* < 0.0001.

Hepatic oxygen consumption (VO₂) was measured continuously to monitor metabolic activity during perfusion. Insulin had no significant effect on VO₂ or on the estimated fluxes, so insulin-treated and untreated livers were pooled within each nutritional state and glucose concentration for all analyses. In the absence of glucose, fasted livers consumed more oxygen than fed livers (Figure 1D). This difference persisted after glucose was added. In Stage 2, nutritional state had a strong effect on VO₂ (*p* < 0.0001), whereas glucose concentration did not (*p* = 0.31; Figure 1E). Nutritional state remained significant in Stage 3 (*p* = 0.017; Figure 1F). Within individual livers, VO₂ did not change significantly across the three stages at either glucose concentration (Figures 1G and 1H). Thus, oxygen consumption was maintained throughout the 90-min protocol despite serial lobectomy, and it was determined by the nutritional state of the donor rather than by perfusate glucose.

### A homogeneous one-compartment model cannot reconcile intracellular isotopomer distributions

Glucose fragment-based models represent the liver as a single, well-mixed compartment. To test this assumption, a one-compartment network comprising glycolysis, gluconeogenesis, glycerol metabolism, glycogen and the TCA cycle was constrained by effluent glucose fragments together with the mass isotopomer distributions (MIDs) of intracellular metabolites (Figure 2A). Each liver contributed two datasets, one at 0 mM glucose (Stage 1) and one at 5 or 20 mM glucose, giving 45 datasets from 23 livers (see Methods). At 5 and 20 mM glucose, the Stage 2 ([U-¹³C₃]lactate/pyruvate) and Stage 3 ([U-¹³C₆]glucose) samples from the same liver were fitted simultaneously as parallel labeling experiments, so that a single flux solution had to account for labeling from both gluconeogenic and glycolytic tracers. With the full set of intracellular metabolites, the model fitted none of 11 fasted and 5 of 12 fed datasets at 0 mM glucose by the χ² test (Figures 2B and 2C). Across all six conditions, only 5 of 45 datasets yielded statistically acceptable fits.

**Figure 2.**
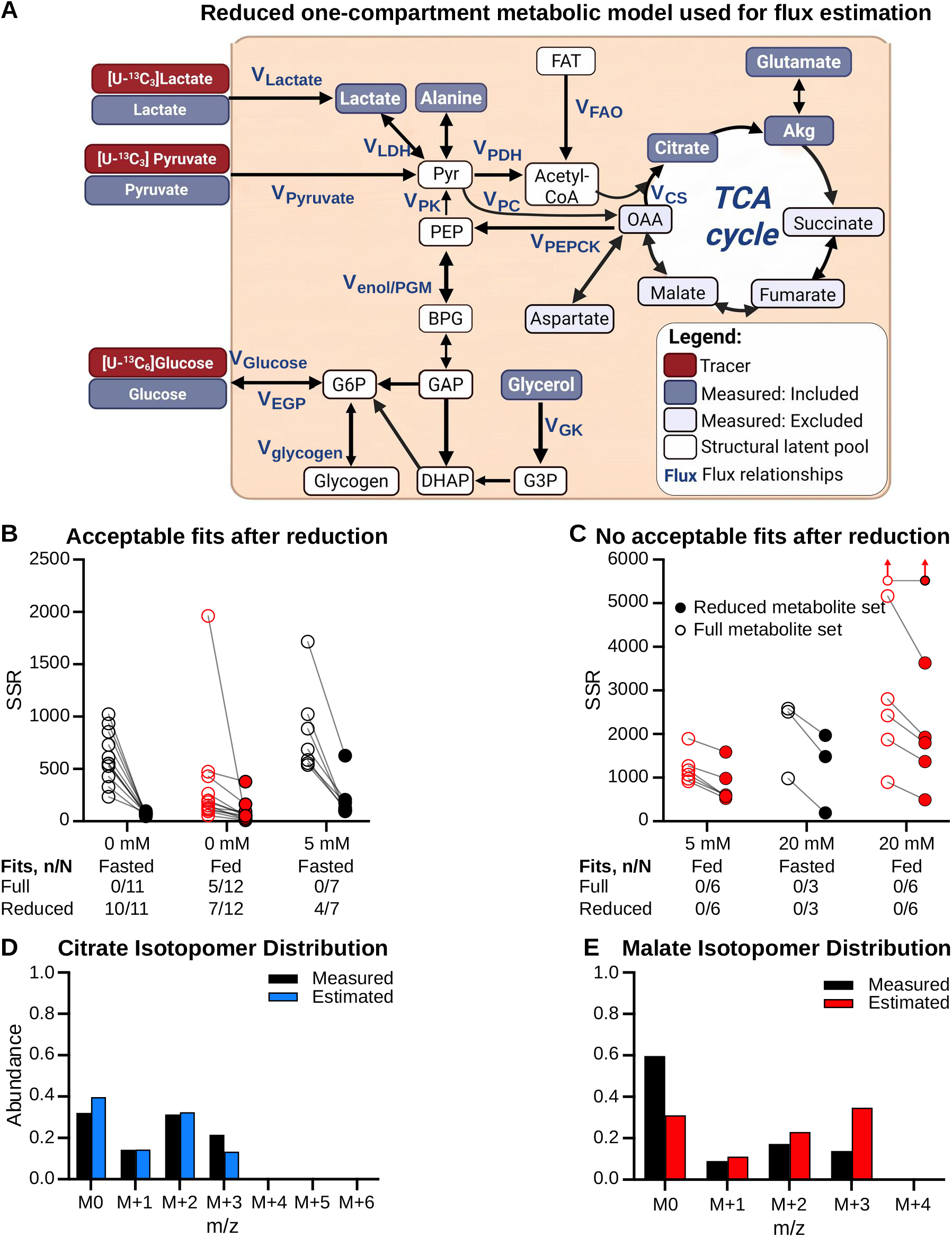
A homogeneous one-compartment model cannot reconcile the full intracellular isotopomer dataset. (A) Reduced one-compartment network used for ^13^C metabolic flux analysis, with tracer inputs from [U-^13^C_3_]lactate, [U-^13^C_3_]pyruvate and [U-^13^C_6_]glucose. Node shading distinguishes tracers, measured metabolites retained as fitting constraints (Measured: included), measured metabolites removed during model reduction (Measured: excluded) and unmeasured structural pools. Modeled fluxes (V) are labeled in blue. In the reduced model, citrate and α-ketoglutarate were the only TCA cycle intermediates retained as constraints. For one fasted liver at 5 mM glucose, intracellular lactate was also excluded to obtain an acceptable fit. (B and C) Sum of squared residuals (SSR) for each liver fitted with the full (open circles) or reduced (filled circles) intracellular metabolite set; lines connect the two fits of the same liver. Black, fasted; red, fed. Conditions are grouped by whether model reduction produced any acceptable fits (B; fasted and fed at 0 mM glucose, fasted at 5 mM) or none (C; fed at 5 mM, fasted and fed at 20 mM). The tables give the number of datasets with an acceptable fit (n/N) by χ^2^ test of the SSR (see Methods). Note the different y-axis ranges in (B) and (C). Upward arrows mark one fed liver at 20 mM whose one-compartment SSR exceeded the axis range (full set, 13,573; reduced set, 10,097). (D and E) Measured (black) and model-estimated (colored) mass isotopomer distributions of citrate (m/z 459; D) and malate (m/z 419; E) generated by the same one-compartment solution, fitted with the full metabolite set, for one representative fasted liver perfused at 0 mM glucose. The same solution reproduces citrate labeling but not malate labeling, indicating a metabolite-specific lack of fit rather than a general failure of optimization. N = datasets subjected to model fitting, one per liver per glucose condition: 11 fasted and 12 fed at 0 mM, 7 fasted and 6 fed at 5 mM, and 3 fasted and 6 fed at 20 mM glucose (45 datasets from 23 livers). The 20 mM datasets of one fasted liver were excluded for technical reasons (see Methods).

Removing intracellular constraints, with the network unchanged, localized the lack of fit to the TCA cycle. Restricting TCA cycle constraints to citrate and α-ketoglutarate, by removing succinate, fumarate, malate and aspartate, yielded acceptable fits in 10 of 11 fasted and 7 of 12 fed datasets at 0 mM glucose, and in 4 of 7 fasted datasets at 5 mM glucose (Figure 2B); for one of these, intracellular lactate was also excluded to obtain an acceptable fit. Even with this reduced data set, no acceptable fits were obtained for fed livers at 5 mM glucose or for any dataset at 20 mM glucose (Figure 2C). Thus, the model failed most consistently in the conditions with the highest exogenous glucose availability.

Inspection of individual MIDs showed that the lack of fit was metabolite-specific. In a representative fasted liver, the one-compartment solution fitted with the full metabolite set reproduced citrate labeling closely but not malate labeling (Figures 2D and 2E). Because both MIDs were generated by the same flux solution, this discrepancy cannot be explained by a general failure of optimization. Instead, it indicates that some intracellular labeling patterns are incompatible with a single homogeneous network.

### Accepted one-compartment solutions recover the expected fed–fasted differences in hepatic flux

Where the reduced one-compartment model could be fitted, its flux estimates were physiologically coherent (Figure 3). This analysis was necessarily limited to the three conditions that yielded acceptable fits: fasted livers at 0 and 5 mM glucose and fed livers at 0 mM glucose. Pyruvate dehydrogenase flux (V_PDH_) was highest in fed livers (Kruskal–Wallis *p* = 0.0003; Figure 3A). Net flux through enolase/phosphoglycerate mutase (V_GnG_) was gluconeogenic in fasted livers and glycolytic in fed livers (*p* = 0.001; Figure 3F). Lactate dehydrogenase flux (V_LDH_) likewise reversed from net lactate → pyruvate in fasted livers to net pyruvate → lactate in fed livers (*p* = 0.0055; Figure 3G). Pyruvate carboxylase and phosphoenolpyruvate carboxykinase flux (V_PC_, V_PEPCK_) differed across conditions (*p* = 0.037; Figures 3D and 3E), although no individual comparison reached significance, whereas β-oxidation (V_βOX_) and citrate synthase (V_CS_) did not differ (Figures 3B and 3C).

**Figure 3.**
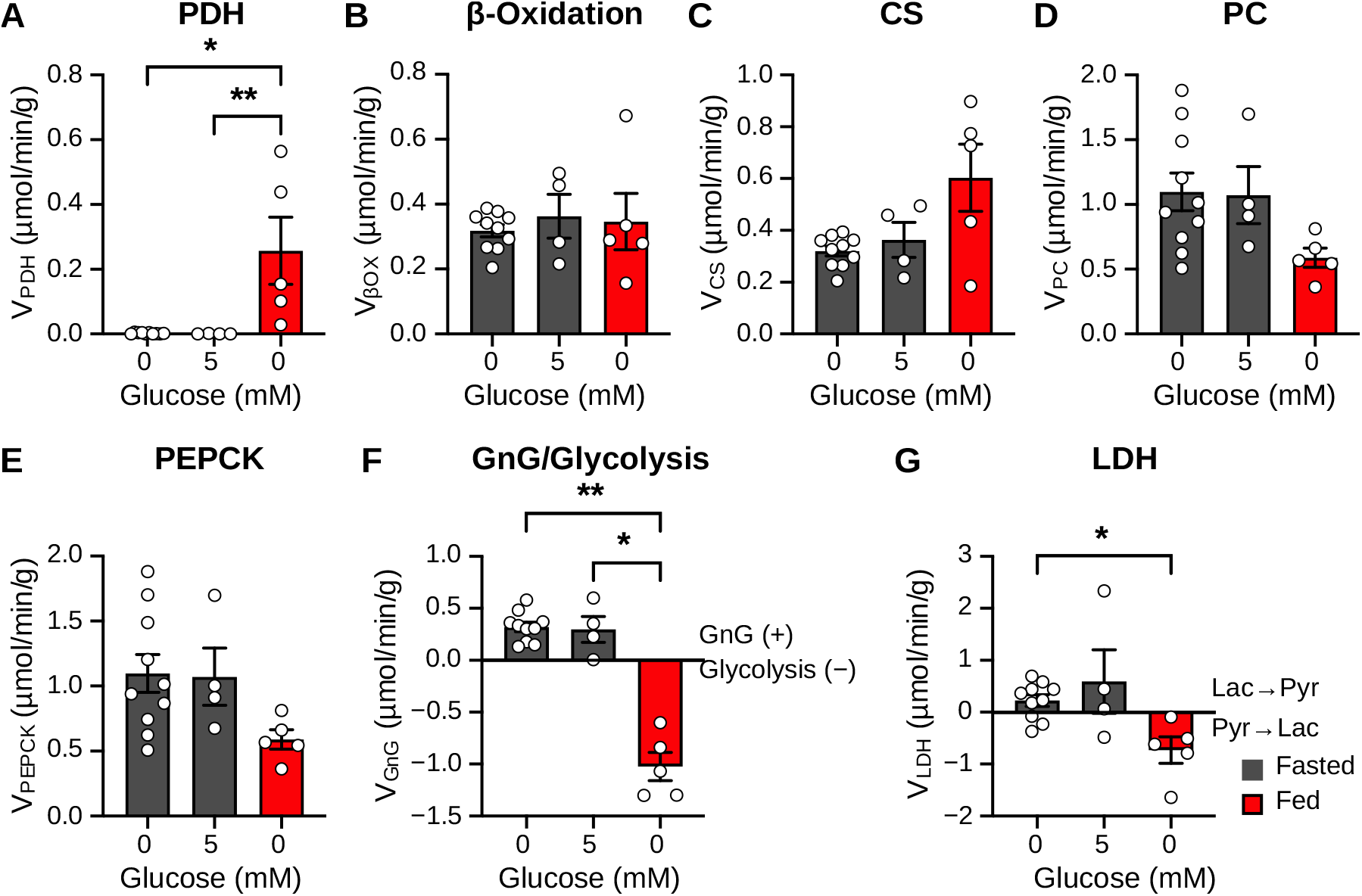
Accepted one-compartment solutions resolve nutritional-state differences in pyruvate oxidation, gluconeogenesis and lactate exchange. Fluxes from reduced one-compartment solutions that passed the χ^2^ goodness-of-fit test (Figure 2B). Only conditions that yielded acceptable fits are shown: fasted livers at 0 and 5 mM glucose and fed livers at 0 mM glucose. (A) Pyruvate dehydrogenase (V_PDH_). (B) β-oxidation (V_βOX_). (C) Citrate synthase (V_CS_). (D) Pyruvate carboxylase (V_PC_). (E) Phosphoenolpyruvate carboxykinase (V_PEPCK_). (F) Net flux through enolase/phosphoglycerate mutase (V_GnG_); positive values denote net gluconeogenic flux and negative values net glycolytic flux. (G) Lactate dehydrogenase (V_LDH_); positive values denote net lactate → pyruvate flux and negative values net pyruvate → lactate flux. V_PC_ and V_PEPCK_ are constrained to be equal by the network structure and are shown separately for comparison with Figure 5. Data are mean ± SEM; each point is one liver. Gray, fasted; red, fed. n = 10 (fasted, 0 mM), 4 (fasted, 5 mM) and 5 (fed, 0 mM) livers with accepted fits. Two of the seven accepted fed 0 mM fits were excluded because their flux estimates reached the parameter bounds of the model. Fluxes are expressed as µmol/min/g liver. Kruskal–Wallis test with Dunn’s multiple-comparisons test. Omnibus *p* values: V_PDH_, 0.0003; V_βOX_, 0.8511; V_CS_, 0.1863; V_PC_, 0.0367; V_PEPCK_, 0.0367; V_GnG_, 0.001; V_LDH_, 0.0055. \**p* < 0.05, \*\**p* < 0.01. See also Figure S1.

Adding 5 mM glucose did not significantly change any flux within the four fasted livers that yielded acceptable fits at both concentrations (Figure S1). With four pairs, however, the smallest attainable *p* value was 0.125, so this comparison cannot exclude a change. Thus, the simplified model recovered the expected nutritional-state differences in pyruvate oxidation, gluconeogenesis and lactate exchange in the three conditions it could fit, but it could not represent the full intracellular dataset.

### A two-compartment model reconciles intracellular and glucose isotopomer data

The metabolite-specific lack of fit indicated that the intracellular data contained labeling from more than one metabolic environment. A second compartment was therefore added, retaining the same biochemical reactions and measurements (Figure 4A). The two compartments were defined by network structure alone and were not assigned to a cell type or lobular zone. The main compartment contained the complete gluconeogenic, glycogenolytic, glycerol, fatty acid oxidation and TCA cycle network. The secondary compartment lacked PC, PEPCK and fatty acid oxidation. It took up glucose from the sinusoidal glucose pool, which contains perfusate glucose and glucose released by the main compartment, and metabolized it through glycolysis. Its pyruvate exchanged with lactate and alanine, which in turn exchanged with the perfusate, or entered the TCA cycle through PDH. Because tissue extracts contain metabolites from both compartments, each measured intracellular MID was modeled as a mixture of the corresponding main-and secondary-compartment pools, with the contribution of each pool estimated during fitting. Effluent lactate and pyruvate likewise integrated output from both compartments, whereas effluent glucose was released only by the main compartment.

**Figure 4.**
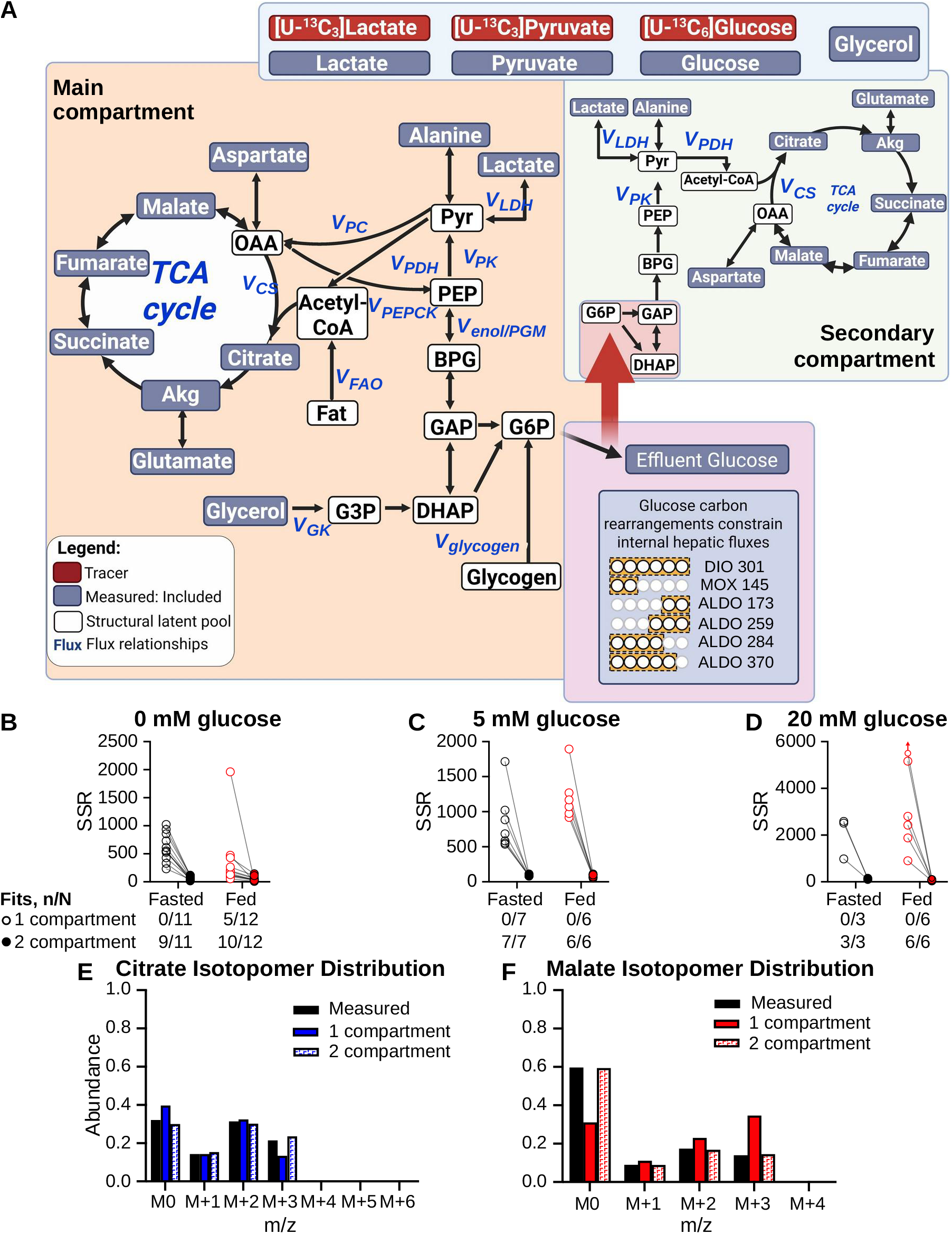
A two-compartment model reconciles intracellular and glucose isotopomer data across nutritional states and glucose concentrations. **(A)** Two-compartment network. Compartments are defined only by network structure and are not assigned to a cell type or lobular zone a priori. The main compartment contains the complete gluconeogenic, glycogenolytic, glycerol, fatty acid oxidation and TCA cycle network. The secondary compartment has no pyruvate carboxylase, phosphoenolpyruvate carboxykinase or fatty acid oxidation: it takes up glucose from the sinusoidal pool (perfusate glucose plus glucose released by the main compartment) and metabolizes it through glycolysis, and its pyruvate exchanges with lactate and alanine or is oxidized through pyruvate dehydrogenase and the TCA cycle. Each measured intracellular metabolite is modeled as a mixture of its main-and secondary-compartment pools. Tracers are [U-^13^C_3_]lactate, [U-^13^C_3_]pyruvate and [U-^13^C_6_]glucose. Node shading follows Figure 2A. Mass isotopomers of effluent glucose fragments provide positional constraints on hepatic fluxes: di-O-isopropylidene propionate m/z 301 (C1–C6), methyloxime pentapropionate m/z 145 (C1–C2) and aldonitrile pentapropionate m/z 173 (C5–C6), 259 (C4–C6), 284 (C1–C4) and 370 (C1–C5). (B–D) SSR for one-compartment (open circles) and two-compartment (filled circles) fits of the same livers at 0 (B), 5 (C) and 20 mM (D) glucose; lines connect the two fits of each liver. Black, fasted; red, fed. One-compartment fits use the full intracellular metabolite set (Figures 2B and 2C). Upward arrow marks one fed liver at 20 mM whose one-compartment SSR (13,573) exceeded the axis range. The tables give the number of datasets with an acceptable fit (n/N) by χ^2^ test of the SSR. Fits were rejected only when the SSR exceeded the upper χ^2^ bound; an SSR below the lower bound was not taken as evidence of lack of fit. The two-compartment model fitted 41 of 45 datasets, compared with 5 of 45 for the one-compartment model. (E and F) Measured (black) mass isotopomer distributions of citrate (m/z 459; E) and malate (m/z 419; F), with distributions predicted by the one-compartment (solid bars) and two-compartment (hatched bars) solutions for the same representative fasted liver at 0 mM glucose as in Figures 2D and 2E. N = datasets subjected to model fitting, as in Figure 2 (45 datasets from 23 livers).

With the full intracellular metabolite set, the two-compartment model yielded acceptable fits in 9 of 11 fasted and 10 of 12 fed datasets at 0 mM glucose and in every dataset at 5 and 20 mM glucose (Figures 4B–4D). Overall, it fitted 41 of 45 datasets, compared with 5 of 45 for the one-compartment model fitted to the same measurements. The improvement was not uniform across metabolites. Citrate labeling, which the one-compartment model already reproduced, remained well fitted, whereas the discordant malate MID was largely reconciled (Figures 4E and 4F). Accepted two-compartment fits had a sum of squared residuals (SSR) within or below the expected χ² range. Values below the lower bound were most frequent at 5 and 20 mM glucose, where unlabeled perfusate glucose diluted ¹³C enrichment and many MIDs therefore carried little label relative to their assigned measurement errors. Such values reflect conservatively assigned errors rather than lack of fit, and are distinct from the excess residual error that defined failure of the one-compartment model.

### Nutritional state is the dominant determinant of main-compartment flux

Across nutritional states and glucose concentrations, main-compartment fluxes were determined mainly by nutritional state, which affected six of the seven fluxes examined; only V_CS_ was unaffected (Figure 5). Fasted livers had higher V_βOX_, V_PC_ and V_PEPCK_ and net gluconeogenic V_GnG_, whereas fed livers had higher V_PDH_, net glycolytic V_GnG_ and net pyruvate → lactate flux (Figures 5A–5G). Glucose concentration had smaller and more selective effects, reaching significance for V_CS_, V_βOX_, V_PC_ and V_PEPCK_ (*p* = 0.028–0.046). Significant interactions for V_βOX_, V_PC_/V_PEPCK_ and V_GnG_ indicated that the response to glucose depended on nutritional state.

**Figure 5.**
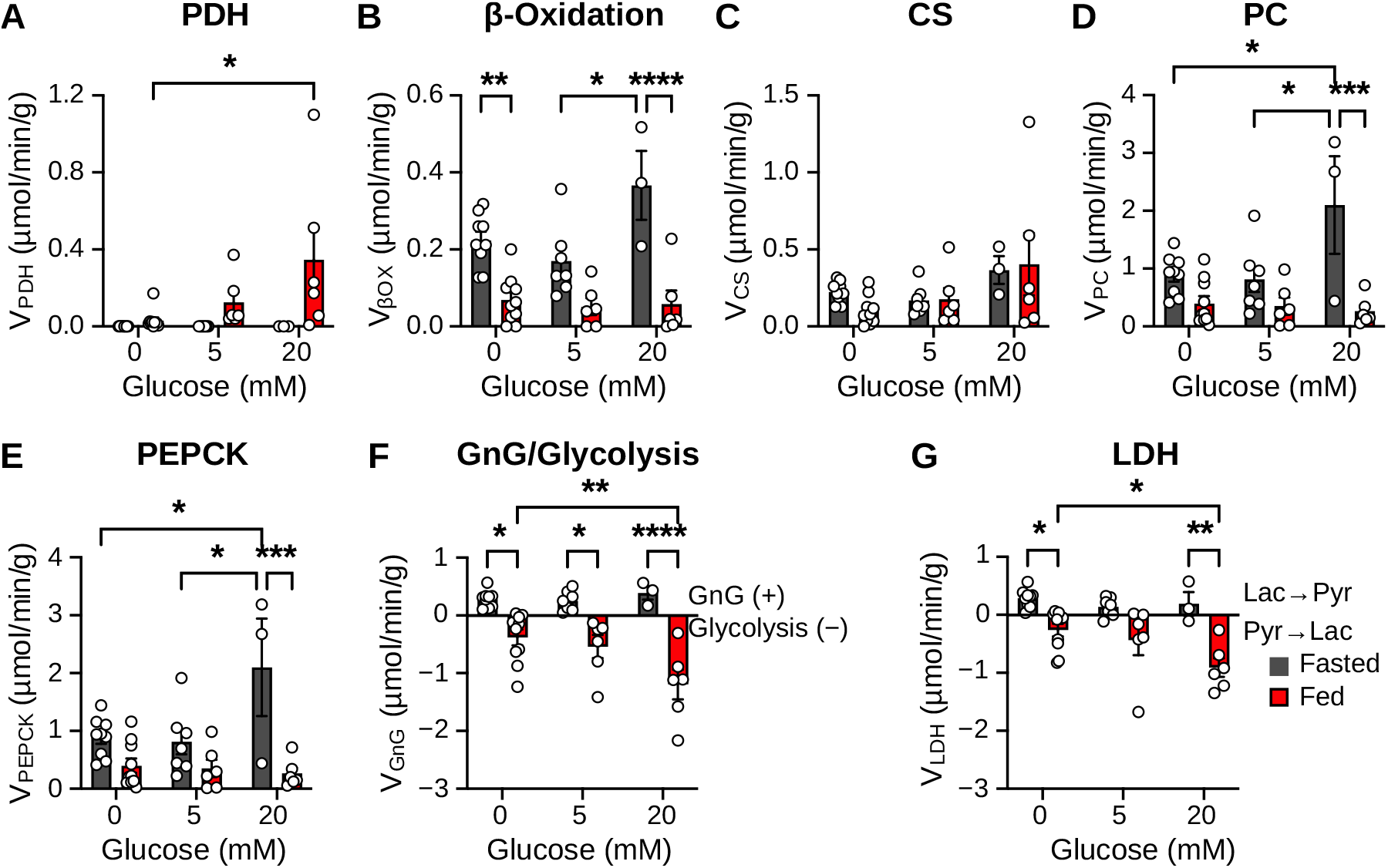
Nutritional state is the dominant determinant of main-compartment fluxes in the two-compartment model. Main-compartment (Figure 4A) fluxes from accepted two-compartment solutions in fasted (gray) and fed (red) livers perfused at 0, 5 and 20 mM glucose. (A) V_PDH_. (B) V_βOX_. (C) V_CS_. (D) V_PC_. (E) V_PEPCK_. (F) V_GnG_; positive values denote net gluconeogenic flux and negative values net glycolytic flux. (G) V_LDH_; positive values denote net lactate → pyruvate flux. V_PC_ and V_PEPCK_ are constrained to be equal by the network structure and are shown separately to parallel Figure 3. Data are mean ± SEM; each point is one liver. n = 9 fasted and 10 fed livers at 0 mM, 7 fasted and 6 fed at 5 mM, and 3 fasted and 6 fed at 20 mM glucose. Fluxes are expressed as µmol/min/g liver. Two-way ANOVA (nutritional state × glucose concentration) with Šídák’s multiple-comparisons test. Nutritional state: V_PDH_, *p* = 0.0048; V_βOX_, V_LDH_, V_PC_, V_PEPCK_ and V_GnG_, each *p* < 0.0001; V_CS_, not significant. Glucose concentration: V_CS_, *p* = 0.0464; V_βOX_, *p* = 0.0279; V_PC_ and V_PEPCK_, *p* = 0.0319; V_PDH_, V_LDH_ and V_GnG_, not significant. Interaction: V_βOX_, *p* = 0.0342; V_PC_ and V_PEPCK_, *p* = 0.0087; V_GnG_, *p* = 0.0252; V_PDH_, V_CS_ and V_LDH_, not significant. \**p* < 0.05, \*\**p* < 0.01, \*\*\**p* < 0.001, \*\*\*\**p* < 0.0001. See also Figures S2 and S3.

Cross-sectional differences between glucose groups were not reproduced within individual livers. In six fasted livers studied at 0 and then 5 mM glucose, no main-compartment flux changed significantly, and individual livers changed in both directions (Figure S2). Together, these data indicate that the metabolic state established before procurement persisted during perfusion and was not overridden by an acute change in glucose supply.

### Flux partitioning between compartments depends on nutritional state

Because main-and secondary-compartment fluxes were estimated simultaneously, the distribution of each pathway between compartments could be compared within the same liver (Figure 6). In fasted livers, all four pathways examined differed significantly between compartments. Net hexose synthesis (V_GHS_) was confined to the main compartment at every glucose concentration (compartment *p* < 0.0001; Figure 6D). V_CS_ was higher in the main compartment at 0 and 20 mM glucose (Figure 6B), and V_LDH_ was net lactate → pyruvate in the main compartment but near zero or net pyruvate → lactate in the secondary compartment (Figure 6C). V_PDH_ was low in both compartments except at 5 mM glucose, when it was higher in the secondary compartment (Figure 6A). Because the secondary compartment lacks fatty acid oxidation, its acetyl-CoA is supplied entirely by pyruvate oxidation.

**Figure 6.**
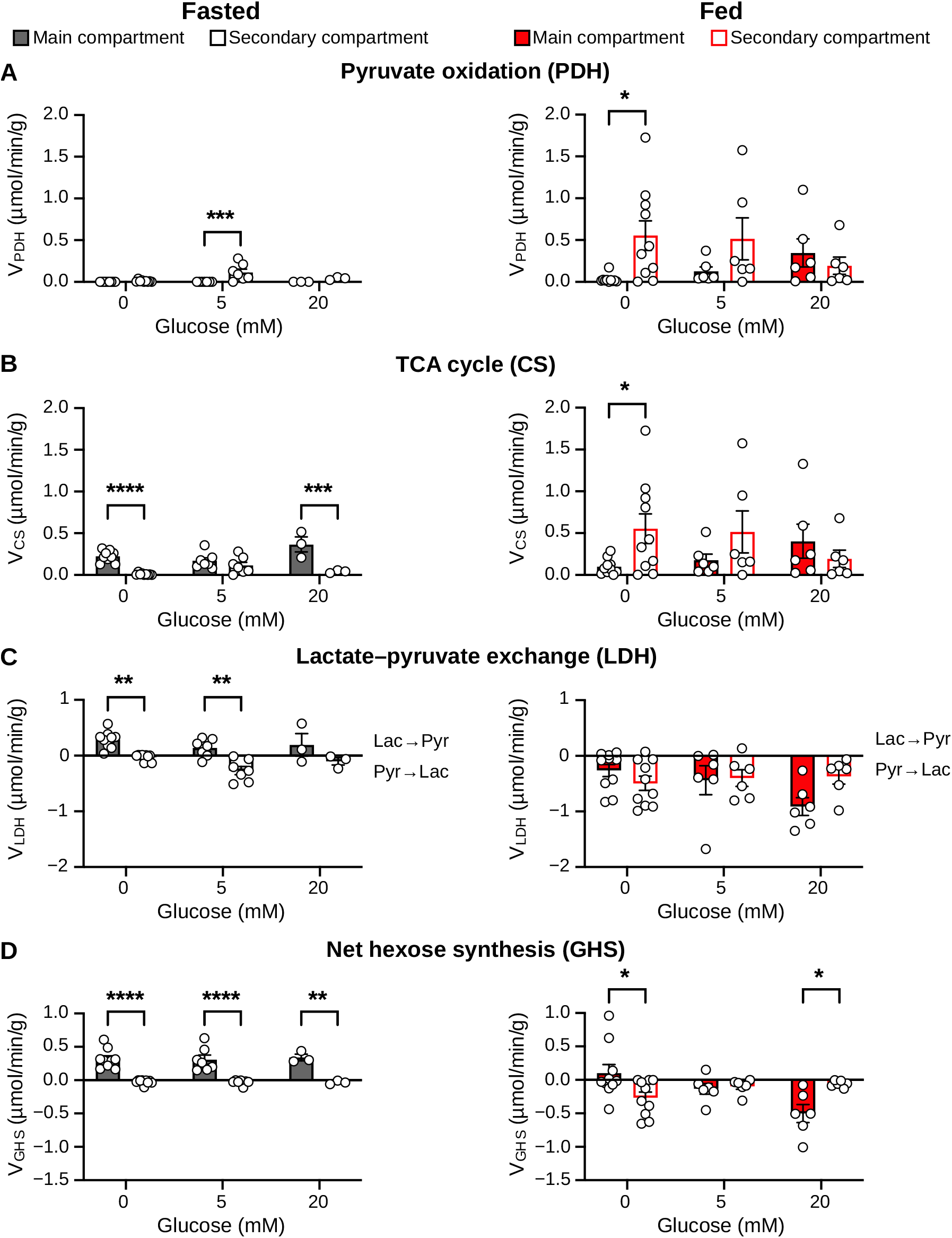
Partitioning of pyruvate oxidation, TCA cycle flux, lactate exchange and hexose synthesis between model compartments. Main-compartment. (filled bars) and secondary-compartment (open bars) fluxes from the same two-compartment solutions in fasted (left; gray) and fed (right; red) livers at 0, 5 and 20 mM glucose. Because the secondary compartment has no fatty acid oxidation, its acetyl-CoA is derived entirely from pyruvate dehydrogenase. (A) Pyruvate oxidation (V_PDH_). (B) TCA cycle flux (V_CS_). (C) Lactate–pyruvate exchange (V_LDH_); positive values denote net lactate → pyruvate flux and negative values net pyruvate → lactate flux. (D) Net hexose synthesis (V_GHS_), the net flux from triose phosphates to glucose-6-phosphate; unlike V_GnG_ (Figures 3 and 5), it includes glycerol-derived carbon. Positive values denote net hexose phosphate synthesis and negative values net glycolytic flux. Fasted and fed panels in each row share the same y-axis range. Data are mean ± SEM; each point is one liver, and main-and secondary-compartment values from the same liver are paired. n = 9, 7 and 3 fasted livers and 10, 6 and 6 fed livers at 0, 5 and 20 mM glucose, respectively. Fluxes are expressed as µmol/min/g liver. Fasted and fed livers were analyzed separately by two-way repeated-measures ANOVA (glucose concentration × compartment) with Šídák’s multiple-comparisons test. Fasted: V_PDH_, glucose *p* = 0.0146, compartment *p* = 0.0032, interaction *p* = 0.0136; V_CS_, compartment *p* < 0.0001, interaction *p* = 0.0095, glucose not significant; V_LDH_, glucose *p* = 0.0012, compartment *p* = 0.0003, interaction not significant; V_GHS_, compartment *p* < 0.0001, glucose and interaction not significant. Fed: no significant main effect or interaction for V_PDH_, V_CS_ or V_LDH_; V_GHS_, interaction *p* = 0.0050, main effects not significant. Šídák’s pairwise comparisons were performed irrespective of the omnibus result; asterisks for fed V_PDH_ and V_CS_ at 0 mM therefore indicate pairwise differences in the absence of a significant overall effect. \**p* < 0.05, \*\**p* < 0.01, \*\*\**p* < 0.001, \*\*\*\**p* < 0.0001.

Compartmental differences were weaker in fed livers. No overall compartment effect was detected for V_PDH_, V_CS_ or V_LDH_. V_GHS_ showed a significant compartment × glucose interaction (*p* = 0.005): net glycolytic flux was located mainly in the secondary compartment at 0 mM glucose but in the main compartment at 20 mM glucose (Figure 6D). Thus, the two compartments were not metabolically interchangeable, and their division of labor depended on nutritional state and, in fed livers, on glucose supply.

### Glucose fragment-based flux estimates track two-compartment estimates

Fluxes estimated by the simplified glucose fragment model used in vivo, which is constrained by effluent glucose labeling alone, tracked main-compartment fluxes estimated by the two-compartment model. In eight fasted livers perfused without glucose, including four whose glucose fragment model fits exceeded the upper χ² bound (Table S3A), estimates from the two approaches were strongly correlated for V_βOX_ (R² = 0.80), V_CS_ (R² = 0.81), V_PC_ (R² = 0.88), V_LDH_ (R² = 0.89) and V_GnG_ (R² = 0.68; Figure S3). Agreement in absolute magnitude was pathway-dependent. Regression slopes were close to 1 for V_βOX_ and V_CS_ (1.11 and 1.10), but lower for V_PC_, V_GnG_ and V_LDH_ (0.59, 0.64 and 0.72). Therefore, the glucose fragment model preserved the ranking of livers for all five fluxes, but gave lower absolute values for the anaplerotic, gluconeogenic and LDH fluxes when these fluxes were high.

## DISCUSSION

By sampling intracellular metabolites during ex vivo perfusion, we tested an assumption that underlies most in vivo estimates of hepatic flux: that the liver can be modeled as a single, homogeneous metabolic compartment. The intracellular data could not be reconciled with this assumption. The lack of fit was reproducible and metabolite-specific, and it was largely resolved by a model with two interconnected compartments of different metabolic capacity. Even so, a simplified glucose fragment model recovered the dominant flux relationships estimated by the two-compartment model. These two observations are compatible: glucose labeling integrates metabolism across the liver, whereas intracellular labeling exposes how that metabolism is distributed.

Adding measurements initially made the model fit worse, not better. Simplified glucose fragment models lump pathways and constrain exchange reactions so that fluxes can be estimated from glucose labeling alone^11,15^. When we added intracellular MIDs while keeping these assumptions, most datasets could not be fitted. This failure was structured: citrate was reproduced by the same solution that failed to reproduce malate. Heterogeneous labeling of TCA cycle intermediates has been reported in perfused rat liver and attributed to zonation, incomplete equilibration between cytosolic and mitochondrial pools, or both^25,26,28^. Our results show that this heterogeneity is large enough to make a homogeneous model statistically untenable once intracellular measurements are included. They also show that a second compartment is sufficient to reconcile it.

The two compartments are operational and are defined by the model architecture. Their biological identity cannot be established from these data. Nevertheless, the architecture that fits the data has a plausible counterpart in lobular zonation. Periportal hepatocytes are specialized for gluconeogenesis and oxidative metabolism, whereas pericentral hepatocytes are more glycolytic^16,17,19,20,22^. In a liver perfused from the portal vein, glucose released upstream could be taken up downstream, as the model’s transfer of glucose from the main to the secondary compartment implies. The failure of the one-compartment model under the conditions most likely to favor hepatic glucose uptake is consistent with this interpretation. Incomplete equilibration between subcellular pools and contributions from non-parenchymal cells are alternative or additional explanations^24,26,27^. Spatial metabolomics or zone-specific genetic models will be needed to distinguish these possibilities^22,32^.

Compartmental flux partitioning varied with nutritional state. In fasted livers, hexose synthesis was confined to the main compartment, and the compartments differed in TCA cycle flux and in the direction of lactate exchange. In fed livers, these differences were smaller, and the site of net glycolysis shifted between compartments with glucose concentration. Some partitioning is imposed by the model, because PC, PEPCK and fatty acid oxidation are absent from the secondary compartment by design. However, the dependence of partitioning on nutritional state and glucose supply is not imposed by the network and is therefore a property of the data.

These findings do not invalidate simplified glucose fragment models; rather, they define what such models measure. Estimates from the glucose fragment model were strongly correlated with two-compartment estimates for β-oxidation, citrate synthase, pyruvate carboxylase, LDH and gluconeogenesis. This supports their use for comparing hepatic flux between physiological states or genotypes, as in vivo studies have done^9,11,12,14^. Correlation did not imply equivalence. Slopes below unity for anaplerotic, gluconeogenic and LDH fluxes indicate that absolute values for these pathways depend on how intracellular structure is represented. We therefore propose a hierarchical view: glucose labeling reports organ-level flux relationships robustly, and intracellular measurements add quantitative and spatial resolution where it is needed.

Nutritional state had a stronger influence than perfusate glucose on oxygen consumption, on main-compartment fluxes and on compartmental partitioning. These differences persisted for 90 min after the liver was removed from the animal, and acute exposure to glucose did not consistently alter flux within individual livers. The paired comparisons were small (four and six livers), so modest acute effects cannot be excluded. Nevertheless, the data indicate that metabolic state at procurement continues to shape hepatic metabolism during perfusion. Possible mechanisms include differences in glycogen and lipid stores, enzyme abundance and covalent regulation. Insulin had no significant additional effect in these experiments. Although insulin was delivered via the portal vein, it was infused at a constant concentration independent of nutritional state, whereas portal insulin in vivo is pulsatile and rises with feeding^33^. The perfusate fatty acid concentration was also fixed, removing the indirect suppression of hepatic glucose production that insulin exerts in vivo through inhibition of adipose lipolysis^34^. The absence of an insulin effect therefore reflects the conditions of isolated perfusion and should not be interpreted as hepatic insulin resistance.

These results have practical implications for NMP. Current viability criteria report whether the liver is metabolically active^2,4,5^. MFA can additionally report how metabolism is organized, including the balance between oxidation, anaplerosis and gluconeogenesis^7,35,36^. The finding that donor nutritional state persists during perfusion suggests that metabolic measurements made on the circuit will partly reflect the condition of the donor. The finding that simplified glucose fragment models track the dominant fluxes suggests that practical, effluent-based flux measurements are feasible during perfusion. Whether such measurements predict graft function will require studies in human livers with clinical outcomes^31^.

This study has limitations. First, the compartments are operational: their architecture was specified rather than inferred, some compartment-specific differences therefore reflect model structure, and the secondary labeling environment may include contributions from non-parenchymal cells, sinusoidal perfusate or pools not at isotopic steady state. Second, the two-compartment model has more free parameters, and its fits were frequently below the expected SSR range, consistent with conservative measurement errors^37–39^. Third, analyses of one-compartment fluxes were restricted to the conditions in which that model could be fitted, and some comparisons were small, particularly the paired within-liver analyses and the 20 mM glucose groups. Fourth, the isolated mouse liver lacks the endocrine, neural and inter-organ inputs present in vivo, and our findings require confirmation in human livers under clinical NMP conditions. Finally, we focused on central carbon metabolism; additional tracers for lipid and amino acid metabolism may reveal further compartmentation^40,41^.

In conclusion, intracellular isotope tracing during ex vivo perfusion shows that hepatic metabolic compartmentation is large enough to constrain whole-liver flux estimation, and that the failure of a homogeneous model is therefore informative about how intact hepatic metabolism is organized. Nevertheless, simplified glucose fragment models recover the dominant physiological fluxes. Together, these findings establish a framework in which effluent glucose labeling provides robust organ-level flux estimates and intracellular labeling resolves how flux is distributed, supporting quantitative metabolic assessment of the liver during machine perfusion.

## ACKNOWLEDGMENTS

The authors acknowledge the UT Southwestern NORC Metabolic Phenotyping and Quantitative Metabolism Cores (NIH P30DK127984). Financial support was provided by NIH R01DK078184 (SCB), R01DK128168 (SCB), an endowment from the Dr. Robert C. and Veronica Atkins Foundation (SCB), and the UT Southwestern Center for Human Nutrition.

## AUTHOR CONTRIBUTIONS

Conceptualization, A.M.A. and S.C.B.; Methodology, A.M.A.; Investigation, A.M.A., J.F., M.M., N.B. and M.I.-V.M.; GC–MS analysis, A.M.A., M.M. and N.B.; Formal analysis and flux modeling, A.M.A. and S.C.B.; Visualization, A.M.A.; Writing – original draft, A.M.A.; Writing – review & editing, A.M.A. and S.C.B.; Supervision, A.M.A. and S.C.B.; Funding acquisition and resources, S.C.B.

## DECLARATION OF INTERESTS

The authors declare no competing interests.

## DECLARATION OF GENERATIVE AI AND AI-ASSISTED TECHNOLOGIES

During the preparation of this work, the authors used Claude (Anthropic) to assist with editing the manuscript text. The authors reviewed and edited all content and take full responsibility for the publication.

## METHODS

### Resource availability

#### Lead contact

Further information and requests for resources and reagents should be directed to and will be fulfilled by the lead contact, Amin M. Amin.

#### Materials availability

This study did not generate new unique reagents.

#### Data and code availability

Metabolic network reactions, measurement and boundary-flux constraints, and goodness-of-fit statistics for each model are provided in Tables S1–S3 (Supplemental Information). Isotopomer data and INCA model files are available from the lead contact upon request. This study did not generate original code.

### Experimental model and study participant details

Male C57BL/6J mice (The Jackson Laboratory) were studied at 14–16 weeks of age. Mice were housed at 22–24°C on a 12-h light–dark cycle (lights on 06:00–18:00) with ad libitum access to standard chow (Teklad Global 16% Protein Diet) and water. Fasted mice were fasted for 16 h overnight in wire-bottom cages to prevent coprophagy. All procedures were approved by the Institutional Animal Care and Use Committee at UT Southwestern Medical Center (protocol APN 2018-102548) and were conducted in accordance with the National Institutes of Health Guide for the Care and Use of Laboratory Animals.

### Method details

#### Ex vivo liver perfusion

Livers were perfused using a single-pass (non-recirculating) system adapted from previously described mouse liver perfusion methods^9,30^. Mice were anesthetized with ketamine/xylazine and heparinized, and the portal vein was cannulated. Perfusion was started immediately at a constant flow of 8 mL/min, the inferior vena cava was transected to allow outflow, and the liver was excised and placed in a temperature-controlled chamber. The perfusate was Krebs–Henseleit buffer containing 3% bovine serum albumin and 0.8 mM of a physiological free fatty acid mixture (22% palmitic, 5% palmitoleic, 3% stearic, 27% oleic, 38% linoleic, 2% γ-linolenic and 3% docosahexaenoic acid). It was equilibrated with 95% O₂/5% CO₂ in a hollow-fiber membrane oxygenator and maintained at 37°C with a circulating water bath.

#### Staged tracer protocol

Each liver was perfused for three consecutive 30-min stages (Figures 1B and 1C). In Stage 1, the perfusate contained [U-¹³C₃]lactate (1.5 mM) and [U-¹³C₃]pyruvate (0.15 mM) without glucose. In Stage 2, unlabeled glucose (5 or 20 mM) was added while the lactate and pyruvate tracers were maintained. In Stage 3, the tracer was switched to [U-¹³C₆]glucose at the same total glucose concentration, with unlabeled lactate and pyruvate at the same concentrations. For each stage, the substrate and tracer mixture was prepared in sterile single-use 60-mL syringes and delivered by syringe driver. Where indicated, insulin was infused continuously throughout all three stages at a final perfusate concentration of 2.583 nM. At the end of each stage, a liver lobe was ligated, excised and freeze-clamped in liquid nitrogen, and effluent was collected for metabolite and isotopomer analysis.

#### Oxygen consumption

Oxygen tension was measured continuously with fiber-optic sensors (PreSens) upstream and downstream of the liver. VO₂ was calculated from the inflow–outflow oxygen difference and the perfusion flow rate, and it was normalized to the mass of liver remaining at each stage after lobectomy (µmol O₂/min/g liver).

#### Effluent glucose concentration and glucose fragment analysis

Effluent glucose was measured enzymatically (Autokit Glucose, Fujifilm Wako). For isotopomer analysis, glucose was converted to its aldonitrile pentapropionate, methyloxime pentapropionate and di-O-isopropylidene propionate derivatives and analyzed by GC–MS, as described^10,11,13^. The fragments used were di-O-isopropylidene propionate m/z 301 (C1–C6), methyloxime pentapropionate m/z 145 (C1–C2) and aldonitrile pentapropionate m/z 173 (C5–C6), 259 (C4–C6), 284 (C1–C4) and 370 (C1–C5).

#### Intracellular metabolite isotopomers

Frozen liver was extracted and derivatized with methoxylamine hydrochloride and MTBSTFA as described^12^. Lactate, pyruvate, alanine, glycerol, citrate, α-ketoglutarate, succinate, fumarate, malate, glutamate and aspartate were analyzed by GC–MS on an Agilent 7890A gas chromatograph coupled to an Agilent 5975C mass selective detector. MIDs were corrected for natural isotope abundance^42^. GC conditions, derivatization and acquisition ranges are given in the Supplemental Methods (Supplemental Information).

#### Metabolic flux analysis

Fluxes were estimated with INCA^43,44^, which uses the elementary metabolite unit framework^45^. Three networks were used, all expressed in absolute units (µmol/min/g liver). (1) The glucose fragment model was constrained by the six effluent glucose fragment MIDs and measured boundary fluxes (glucose input and output, glycerol input and output, and lactate and pyruvate inputs), without intracellular MIDs, following established in vivo models^11,12^. (2) The one-compartment model used the same reactions but was additionally constrained by intracellular MIDs (Figure 2A). In addition to the glucose fragments and boundary fluxes, the full set comprised intracellular lactate, alanine, succinate, fumarate, α-ketoglutarate, malate, aspartate, citrate and glutamate, together with perfusate and effluent lactate and pyruvate. The reduced set removed succinate, fumarate, malate and aspartate. For one fasted liver at 5 mM glucose, intracellular lactate was also excluded to obtain an acceptable fit. (3) The two-compartment model divided the network into a main compartment, containing the complete network, and a secondary compartment lacking PC, PEPCK, fatty acid oxidation, glycerol metabolism and glycogen, which took up glucose from the sinusoidal pool (perfusate glucose plus glucose released by the main compartment) and exchanged lactate and pyruvate with the perfusate (Figure 4A). The secondary compartment contained a reversible malic enzyme reaction. Each measured intracellular MID was assigned to a mixture of the corresponding main-and secondary-compartment pools, with mixing fractions estimated as free parameters. Unlabeled dilution pools were included for alanine, aspartate and glutamate, together with unlabeled inputs to secondary-compartment aspartate and glutamate. Effluent lactate and pyruvate received output from both compartments, whereas effluent glucose derived only from the main compartment. The model was fitted with the full intracellular metabolite set (Tables S1C and S2D). V_GnG_ denotes net flux through enolase/phosphoglycerate mutase, V_GHS_ denotes net flux from triose phosphates to glucose-6-phosphate, which, unlike V_GnG_, includes glycerol-derived carbon, and V_LDH_ is positive for net lactate → pyruvate flux. Because PC is the only anaplerotic entry and PEPCK the only cataplerotic exit of the main-compartment TCA cycle, V_PC_ = V_PEPCK_ at metabolic steady state in both networks. Datasets at 0 mM glucose comprised Stage 1 measurements. At 5 and 20 mM glucose, Stage 2 and Stage 3 measurements from the same liver were fitted simultaneously in INCA as parallel labeling experiments sharing a single set of fluxes, with each experiment defined by its own tracer inputs. This approximately doubled the number of independent measurements constraining each solution. Reactions and measurements for each model are listed in Tables S1 and S2.

#### Model fitting and goodness of fit

Fluxes were estimated for each liver by weighted least-squares regression, minimizing the SSR between measured and simulated MIDs^37,40^. Each model was fitted with 50 restarts from random initial values perturbed over four orders of magnitude (relative convergence tolerance 0.001; Levenberg–Marquardt τ = 10⁻⁶), and the solution with the lowest SSR was retained. A measurement standard deviation of 0.007 (0.7 mol%) was assigned to all glucose fragment and intracellular MIDs. Goodness of fit was assessed by a χ² test at 95% confidence with degrees of freedom equal to the number of independent measurements minus the number of free parameters^37^. A fit was rejected when the SSR exceeded the upper bound. An SSR below the lower bound was not taken as evidence of lack of fit. Accepted solutions in which flux estimates reached the parameter bounds of the model were excluded from flux comparisons (two fed datasets at 0 mM glucose in the one-compartment model). Confidence intervals were determined by parameter continuation^37^.

### Quantification and statistical analysis

Statistical analyses were performed in GraphPad Prism 10. Data are presented as mean ± SEM. For VO₂, n refers to individual perfused livers. For flux analyses, n refers to datasets: each liver contributed one dataset at 0 mM glucose (Stage 1) and one at its assigned glucose concentration, giving 45 datasets from 23 livers. The Stage 2 and Stage 3 isotopomer data from one fasted liver in the 20 mM group were excluded from flux analyses because the measured labeling patterns were not physiologically interpretable, consistent with sample contamination or mix-up during GC–MS analysis that could not be verified retrospectively. For transparency, fits of these data with both models are reported in Table S3. Its Stage 1 data and its oxygen consumption data were retained. VO₂ in Stage 1 was compared by Welch’s t test. VO₂ in Stages 2 and 3 was analyzed by two-way ANOVA (nutritional state × glucose concentration) with uncorrected Fisher’s LSD test. Within-liver VO₂ trajectories were analyzed by two-way repeated-measures ANOVA with Geisser–Greenhouse correction and Šídák’s test. Fluxes from accepted one-compartment solutions were compared by Kruskal–Wallis test with Dunn’s test. Main-compartment fluxes were analyzed by two-way ANOVA (nutritional state × glucose concentration) with Šídák’s test. Main-and secondary-compartment fluxes were compared within fasted and fed livers by two-way repeated-measures ANOVA (glucose concentration × compartment) with Šídák’s test. Paired within-liver fluxes at 0 and 5 mM glucose were compared by two-sided Wilcoxon matched-pairs signed-rank test. Agreement between models was assessed by least-squares linear regression. The effect of insulin was tested before pooling by two-way ANOVA within each nutritional state and glucose concentration, and no significant insulin effects were detected; insulin-treated and untreated livers were therefore pooled for all analyses. Because each insulin subgroup contained few livers, modest insulin effects cannot be excluded. *p* < 0.05 was considered significant. Statistical details for each experiment are given in the figure legends.

Schematics were created with BioRender.com.

## SUPPLEMENTAL FIGURES

**Figure S1.**
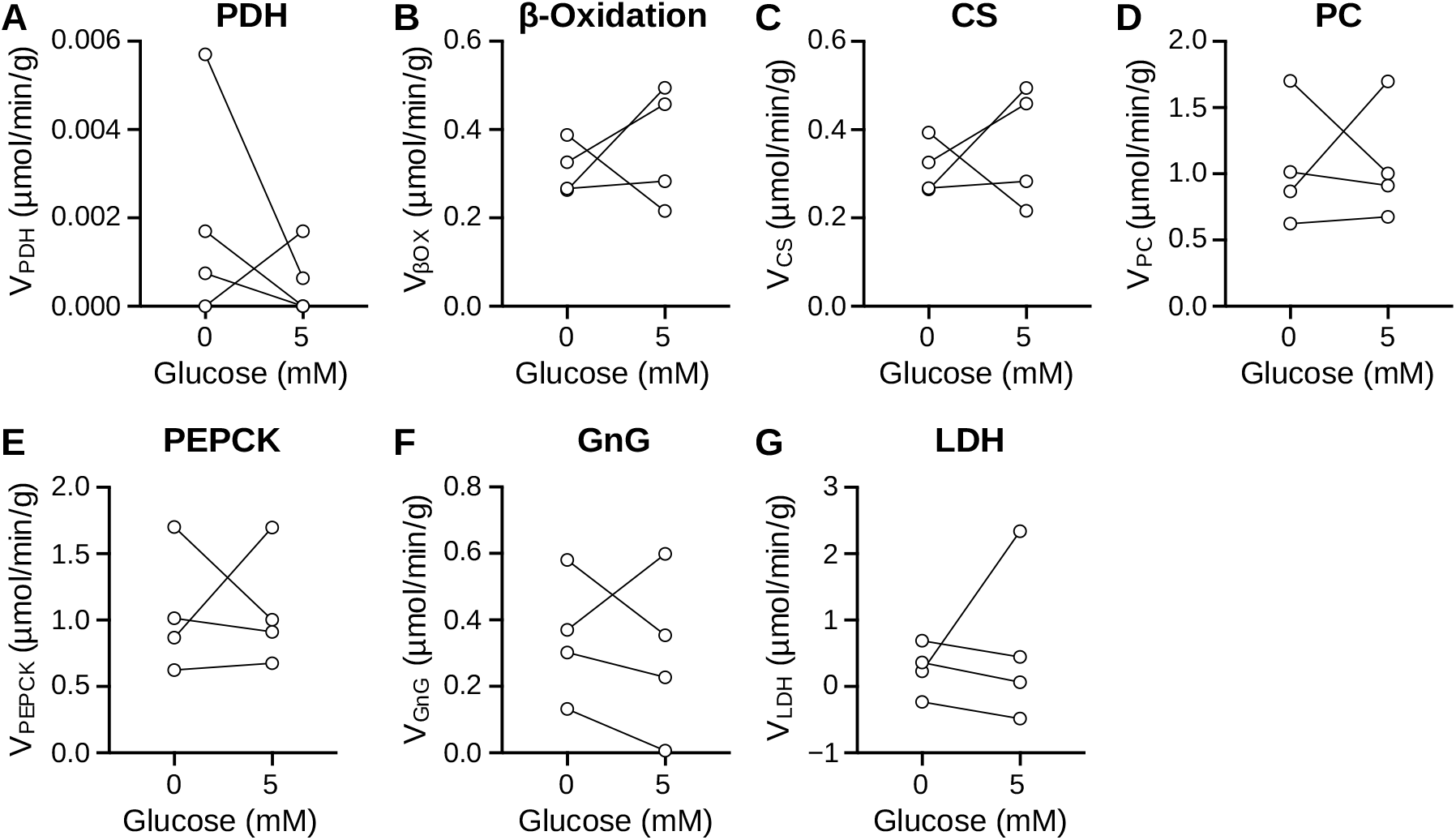
Within-liver one-compartment flux estimates before and after glucose addition, related to. Figure 3. (A–G) Fluxes estimated by the reduced one-compartment model in the same fasted livers at 0 mM glucose and after the transition to 5 mM glucose: V_PDH_ (A), V_βOX_ (B), V_CS_ (C), V_PC_ (D), V_PEPCK_ (E), V_GnG_ (F) and V_LDH_ (G). Lines connect estimates from the same liver. Only livers with accepted one-compartment fits at both glucose concentrations were included. V_PC_ and V_PEPCK_ are equal by network construction. n = 4 fasted livers. Fluxes are expressed as µmol/min/g liver. Two-sided Wilcoxon matched-pairs signed-rank test; exact *p* values: V_PDH_, 0.5000; V_βOX_, 0.6250; V_CS_, 0.6250; V_PC_, >0.9999; V_PEPCK_, >0.9999; V_GnG_, 0.8750; V_LDH_, 0.8750. With four pairs, the smallest attainable two-sided exact *p* value is 0.125, so these comparisons cannot exclude a change in flux.

**Figure S2.**
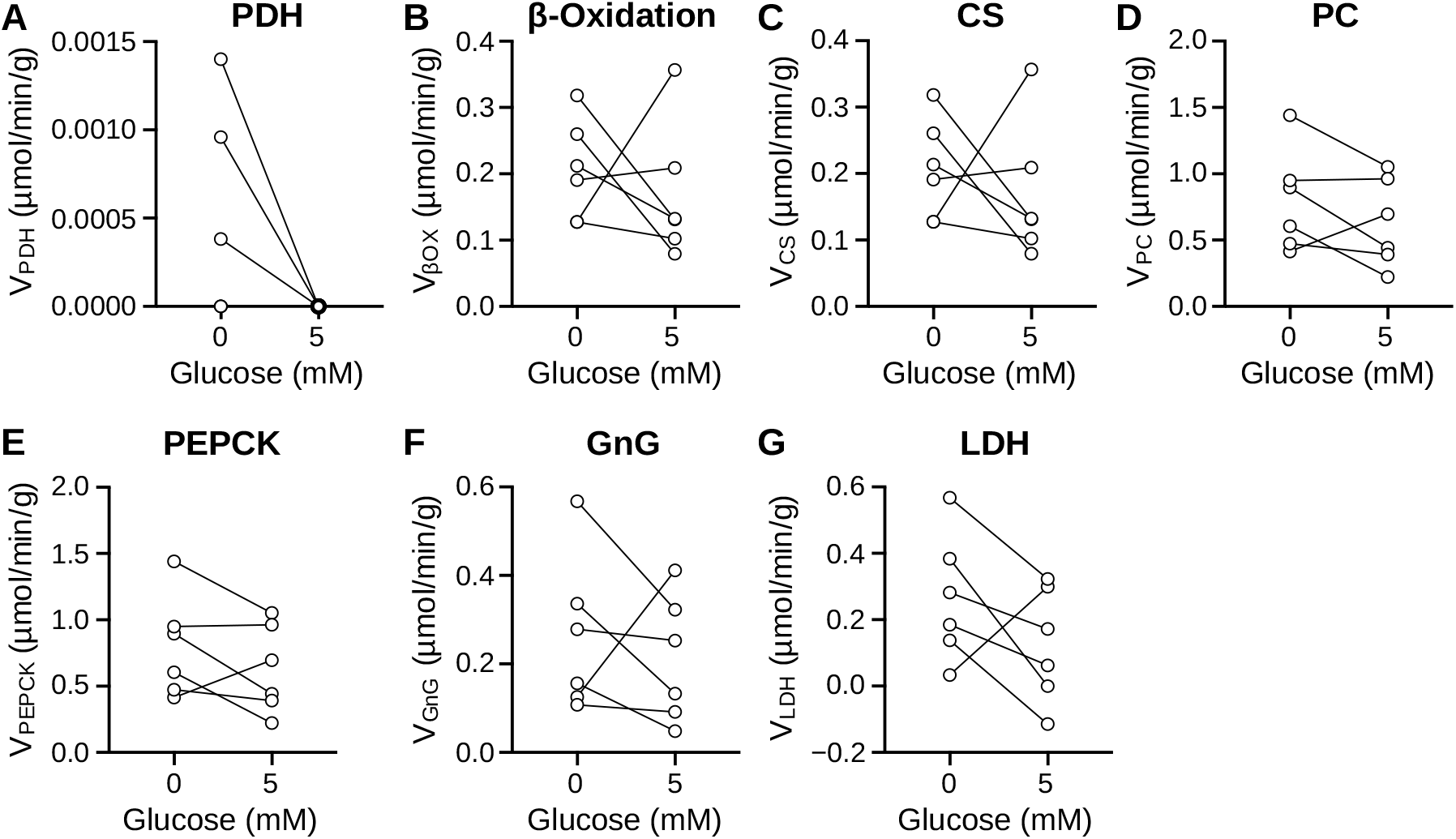
Within-liver main-compartment flux estimates before and after glucose addition, related to. Figure 5. (A–G) Main-compartment fluxes estimated by the two-compartment model in the same fasted livers at 0 mM glucose and after the transition to 5 mM glucose: V_PDH_ (A), V_βOX_ (B), V_CS_ (C), V_PC_ (D), V_PEPCK_ (E), V_GnG_ (F) and V_LDH_ (G). Lines connect estimates from the same liver. Only livers with accepted two-compartment fits at both glucose concentrations were included. n = 6 fasted livers. Fluxes are expressed as µmol/min/g liver. Two-sided Wilcoxon matched-pairs signed-rank test; exact *p* values: V_PDH_, 0.2500; V_βOX_, 0.5625; V_CS_, 0.5625; V_PC_, 0.2188; V_PEPCK_, 0.2188; V_GnG_, 0.4375; V_LDH_, 0.3125. For V_PDH_, three livers showed no change and were excluded from ranking by the test. No flux differed significantly between 0 and 5 mM glucose.

**Figure S3.**
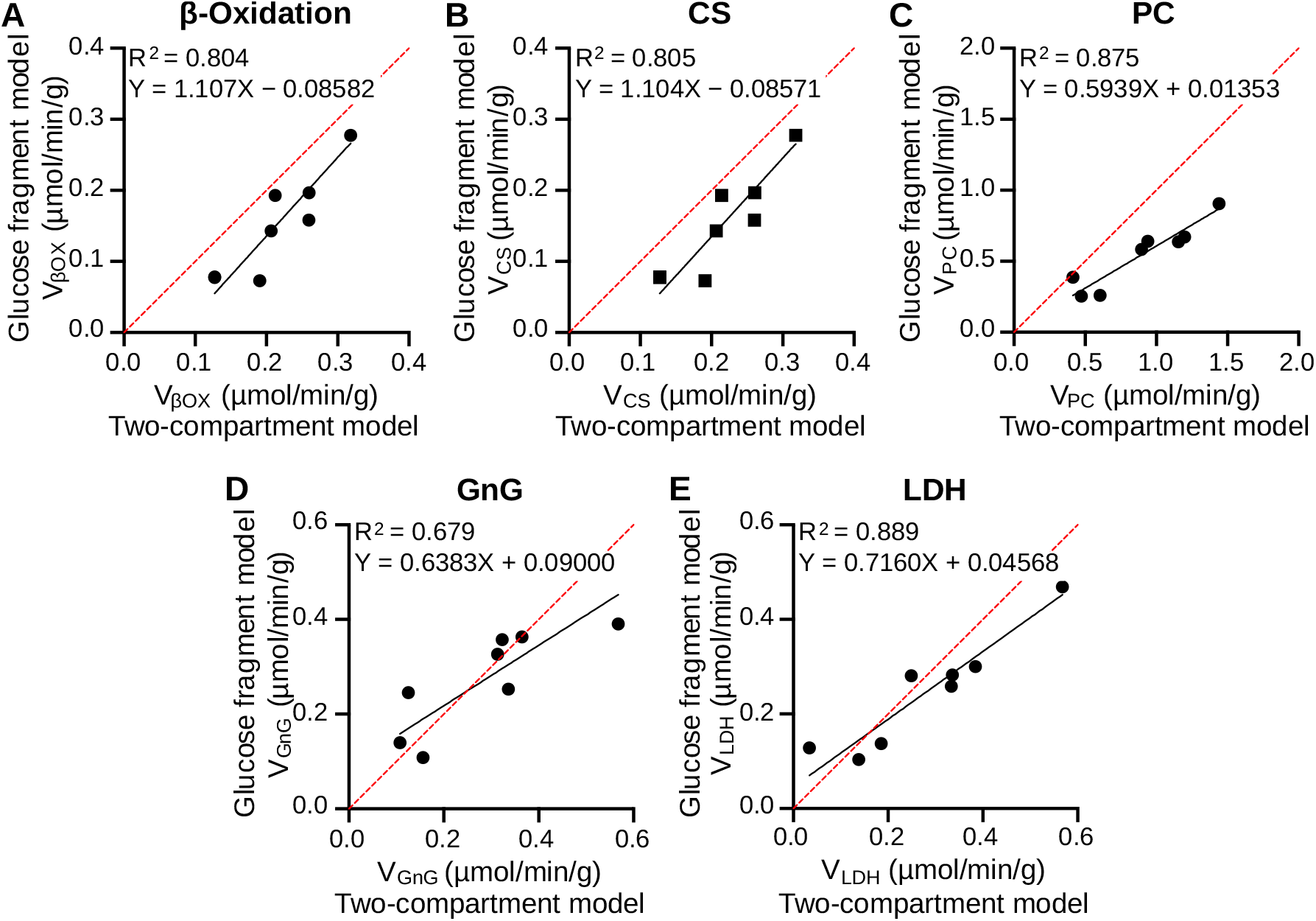
Two-compartment flux estimates correlate with those from a simplified glucose fragment model, related to. Figure 5. (A–E) Fluxes estimated in the same livers by the simplified glucose fragment model used for in vivo studies (y-axis) and by the two-compartment model (main compartment; x-axis): V_βOX_ (A), V_CS_ (B), V_PC_ (C), V_GnG_ (D) and V_LDH_ (E). The red dashed line is the line of identity (y = x); the solid black line is the least-squares linear regression, with its equation and coefficient of determination (R^2^) shown in each panel. n = 8 fasted livers perfused at 0 mM glucose; each point is one liver. Fluxes are expressed as µmol/min/g liver. Glucose fragment model fits for four of these livers exceeded the upper χ^2^ bound (two narrowly; Table S3A) and were retained, because the comparison concerns agreement between flux estimates rather than goodness of fit.

## Notes

### Competing Interest Statement

The authors have declared no competing interest.

