## Supplemental Information for "Resolving Hepatic Metabolic Flux During Ex Vivo Liver Perfusion Reveals Compartment-Specific Constraints"

#### Supplemental Methods

##### Organic acid extraction from liver tissue

Liver tissue was kept frozen until extraction. Approximately 30–50 mg of tissue was weighed into 2-mL microcentrifuge tubes and extracted with 600 µL of prechilled 80:20 (v/v) methanol:water. Samples were thoroughly homogenized on ice, incubated on ice for 10 min, and centrifuged at maximum speed for 10 min at 4°C. The resulting supernatant was transferred to a glass vial and evaporated to complete dryness under a constant stream of nitrogen gas before derivatization.

##### Organic acid extraction from perfusate

To 200 µL of perfusate, 1,000 µL of ice-cold acetone was added. Samples were vortexed thoroughly and centrifuged at maximum speed for 15 min at 4°C. The supernatant was collected and evaporated to complete dryness under nitrogen gas before derivatization.

##### Methoxylamine/MTBSTFA derivatization of organic acids

Following their respective extraction procedures, dried liver and perfusate organic-acid extracts were derivatized using methoxylamine hydrochloride followed by silylation with N-methyl-N-(tert-butyldimethylsilyl)trifluoroacetamide (MTBSTFA). Dried extracts were resuspended in 50 µL of 1% (wt/vol) methoxylamine hydrochloride in pyridine and incubated for 90 min at 37°C. Subsequently, 80 µL of MTBSTFA was added, samples were vortexed, and derivatization was continued for 60 min at 60°C. Samples were then centrifuged at maximum speed for 10 min at room temperature, and 100 µL of supernatant was transferred to a GC–MS vial containing a glass insert. Derivatized samples were analyzed within 24 h.

**Glucose derivatization for positional isotopomer analysis**

Effluent glucose was analyzed separately from organic acids using three complementary glucose derivatives: methyloxime pentapropionate (MOX), aldonitrile pentapropionate (ALDO), and di-O-isopropylidene propionate (DIO). These derivatives generate overlapping fragments containing different subsets of the six glucose carbon atoms, thereby providing positional labeling information for metabolic flux analysis. The six glucose fragment ions incorporated as isotopomer constraints were DIO m/z 301, MOX m/z 145, and ALDO m/z 173, 259, 284, and 370.

**GC–MS instrumentation and acquisition**

GC–MS analysis was performed using an Agilent 7890A gas chromatograph coupled to an Agilent 5975C mass spectrometer equipped with a 70-eV electron-ionization source. Chromatographic separation was performed using an HP-5ms capillary column (30 m × 0.25 mm internal diameter, 0.25-µm film thickness; Agilent J&W). Samples were injected at a volume of 1 µL, with split conditions adjusted to optimize signal-to-noise ratio.

*Glucose derivatives*

For the DIO and ALDO glucose derivatives, the GC oven was initially maintained at 80°C for 1 min, increased at 20°C/min to 280°C, held at 280°C for 4 min, and subsequently increased at 40°C/min to 325°C. For the MOX glucose derivative, the same temperature program was used except that the oven temperature was increased from 80°C to 280°C at 10°C/min. MS data acquisition was initiated after a 5-min solvent delay. Acquisition ranges were m/z 144–150 for MOX, m/z 170–380 for ALDO, and m/z 300–320 for DIO derivatives. Each sample was injected three times.

*Organic acids*

For organic-acid analysis, the GC oven was initially maintained at 150°C for 3 min, increased at 5°C/min to 205°C, increased at 50°C/min to 250°C and held for 1 min, and subsequently increased at 25°C/min to 275°C and held for 3 min. Mass-isotopomer acquisition ranges were m/z 261.1–264.1 for lactate, m/z 274.1–277.1 for pyruvate, m/z 289.1–293.1 for succinate, m/z 287.2–291.2 for fumarate, m/z 419.2–423.2 for malate, m/z 432.2–436.2 for oxaloacetate, m/z 446.2–451.2 for α-ketoglutarate, and m/z 459.2–465.2 for citrate.

**Isotopomer constraints and fragment definitions**

For the final two-compartment metabolic flux model, intracellular isotopomer constraints included lactate m/z 233 and 261, alanine m/z 232 and 260, succinate m/z 289, fumarate m/z 287, α-ketoglutarate m/z 346, malate m/z 391 and 419, aspartate m/z 302, 390, and 418, citrate m/z 459, and glutamate m/z 330 and 432. Extracellular isotopomer constraints included perfusate lactate m/z 233 and 261, effluent lactate m/z 233 and 261, effluent pyruvate m/z 174, and the six effluent glucose fragments DIO m/z 301, MOX m/z 145, and ALDO m/z 173, 259, 284, and 370.

Fragment carbon definitions were specified in INCA according to the metabolite carbon atoms represented by each derivatized ion. For intracellular organic-acid and amino-acid measurements, fragment definitions included lactate m/z 233, C2–C3; lactate m/z 261, C1–C3; alanine m/z 232, C2–C3; alanine m/z 260, C1–C3; succinate m/z 289, C1–C4; fumarate m/z 287, C1–C4; α-ketoglutarate m/z 346, C1–C5; malate m/z 391, C2–C4; malate m/z 419, C1–C4; aspartate m/z 302, C1–C2; aspartate m/z 390, C2–C4; aspartate m/z 418, C1–C4; citrate m/z 459, C1–C6; glutamate m/z 330, C2–C5; and glutamate m/z 432, C1–C5. Corresponding carbon definitions were applied to perfusate and effluent fragments where applicable. For glucose, the modeled carbon definitions were DIO m/z 301, C1–C6; MOX m/z 145, C1–C2; ALDO m/z 173, C5–C6; ALDO m/z 259, C4–C6; ALDO m/z 284, C1–C4; and ALDO m/z 370, C1–C5.

Fragments that were measured but not used as constraints in a given model are identified in Table S2. Fragment availability differed among individual datasets when measurements were unavailable or did not meet analytical quality criteria.

**Mass isotopomer processing and flux analysis**

Mass isotopomer distributions (MIDs) were determined from the relative abundance of each monitored fragment ion and its corresponding mass isotopologues. Measured MIDs were corrected for contributions from naturally occurring isotopes before incorporation into metabolic flux analysis. Corrected isotopomer measurements were supplied to INCA v2.1 as experimental constraints with a measurement standard deviation of 0.007 (0.7 mol%) for all MIDs.

Flux analysis used three networks of increasing intracellular complexity: a glucose fragment model, a one-compartment model and a two-compartment model. At 5 and 20 mM glucose, Stage 2 ([U-$^{13}C_3$]lactate/pyruvate) and Stage 3 ([U-$^{13}C_6$]glucose) measurements from the same liver were fitted simultaneously as parallel labeling experiments sharing a single set of fluxes. Reaction networks are given in Table S1, measurement and boundary-flux constraints in Table S2, and goodness-of-fit statistics in Table S3.

### Table S1. Metabolic network reactions

*Reactions are listed with their INCA identifiers. Pathway categories are given for orientation only. NA, natural abundance.*

#### S1A. Glucose fragment model

| Order | Pathway category | Reaction label | Reaction equation |
|---|---|---|---|
| 1 | Perfusate inputs / outputs | 1 Lac tracer IN | Lac.tracer -> Lac + 0.1*PyrTr |
| 2 | Perfusate inputs / outputs | 1 Pyr tracer IN | Pyr.tracer + PyrTr -> Pyr |
| 3 | Perfusate inputs / outputs | 1 Glc balance | Glc -> Glc.aff |
| 4 | Perfusate inputs / outputs | 1 Glc effluent | Glc.aff -> Glc.eff |
| 5 | Perfusate inputs / outputs | 1 Total Glc OUT | Glc.eff -> Glc.sink |
| 6 | Perfusate inputs / outputs | Bicarb source | Bicarb.source -> CO2.aff |
| 7 | Perfusate inputs / outputs | Bicarb Sink | CO2.aff -> Bicarb.sink |
| 8 | β-oxidation / acetyl-CoA | A1 | Fat -> AcCoA |
| 9 | TCA cycle / anaplerosis / $CO_2$ | A3 | Oac + AcCoA -> Cit |
| 10 | TCA cycle / anaplerosis / $CO_2$ | A4 | Pyr + CO2 -> Oac |
| 11 | TCA cycle / anaplerosis / $CO_2$ | A5 net | Cit <-> Akg + CO2 |
| 12 | TCA cycle / anaplerosis / $CO_2$ | A5 exch | Cit <-> Akg + CO2 |
| 13 | TCA cycle / anaplerosis / $CO_2$ | A6 | Akg -> SucCoA + CO2 |
| 14 | TCA cycle / anaplerosis / $CO_2$ | A8 | SucCoA -> Suc |
| 15 | TCA cycle / anaplerosis / $CO_2$ | A9 net | Suc <-> Fum |
| 16 | TCA cycle / anaplerosis / $CO_2$ | A9 exch | Suc <-> Fum |
| 17 | TCA cycle / anaplerosis / $CO_2$ | A10 net | Fum <-> Mal |
| 18 | TCA cycle / anaplerosis / $CO_2$ | A10 exch | Fum <-> Mal |
| 19 | TCA cycle / anaplerosis / $CO_2$ | A11 net | Mal <-> Oac |
| 20 | TCA cycle / anaplerosis / $CO_2$ | A11 exch | Mal <-> Oac |
| 21 | TCA cycle / anaplerosis / $CO_2$ | T5 net | CO2.aff <-> CO2 |
| 22 | TCA cycle / anaplerosis / $CO_2$ | T5 exch | CO2.aff <-> CO2 |
| 23 | Pyruvate / lactate / alanine exchange | B18 net | Lac <-> Pyr |
| 24 | Pyruvate / lactate / alanine exchange | B18 exch | Lac <-> Pyr |
| 25 | Gluconeogenesis / glycolysis / glycogen | A27* | Oac -> PEP + CO2 |
| 26 | Gluconeogenesis / glycolysis / glycogen | B28 | PEP -> Pyr |
| 27 | Gluconeogenesis / glycolysis / glycogen | B29 | PEP -> BPG |
| 28 | Gluconeogenesis / glycolysis / glycogen | B30 | BPG -> GAP |
| 29 | Gluconeogenesis / glycolysis / glycogen | B31 | Glycerol.aff -> G3P |
| 30 | Gluconeogenesis / glycolysis / glycogen | B32 | G3P -> DHAP |
| 31 | Gluconeogenesis / glycolysis / glycogen | B33 net | DHAP <-> GAP |
| 32 | Gluconeogenesis / glycolysis / glycogen | B33 exch | DHAP <-> GAP |
| 33 | Gluconeogenesis / glycolysis / glycogen | B34 | DHAP + GAP -> G6P |
| 34 | Gluconeogenesis / glycolysis / glycogen | B35 | G6P -> Glc |
| 35 | Gluconeogenesis / glycolysis / glycogen | B36 | Glycogen -> G6P |

#### S1B. One-compartment model

| Order | Pathway category | Reaction label | Reaction equation |
|---|---|---|---|
| 1 | Perfusate inputs / outputs | 1 Lac tracer IN | Lac.tracer -> Lac.perf + 0.1*PyrTr |
| 2 | Perfusate inputs / outputs | 1 NA Lac IN | Lac.source -> Lac.perf + 0.1*PyrNA |
| 3 | Perfusate inputs / outputs | 1 Total Lac IN | Lac.perf -> Lac.aff |
| 4 | Perfusate inputs / outputs | 1 Lac Balance net | Lac.aff <-> Lac |
| 5 | Perfusate inputs / outputs | 1 Lac Balance exch | Lac.aff <-> Lac |
| 6 | Perfusate inputs / outputs | 1 Lac effluent | Lac.aff -> Lac.eff |
| 7 | Perfusate inputs / outputs | 1 Total Lac OUT | Lac.eff -> Lac.sink |
| 8 | Perfusate inputs / outputs | 1 Pyr tracer IN | Pyr.tracer + PyrTr -> Pyr.perf |
| 9 | Perfusate inputs / outputs | 1 NA Pyr IN | Pyr.source + PyrNA -> Pyr.perf |
| 10 | Perfusate inputs / outputs | 1 Total Pyr IN | Pyr.perf -> Pyr.aff |
| 11 | Perfusate inputs / outputs | 1 Pyr Balance net | Pyr.aff <-> Pyr |
| 12 | Perfusate inputs / outputs | 1 Pyr Balance exch | Pyr.aff <-> Pyr |
| 13 | Perfusate inputs / outputs | 1 Pyr effluent | Pyr.aff -> Pyr.eff |
| 14 | Perfusate inputs / outputs | 1 Total Pyr OUT | Pyr.eff -> Pyr.sink |
| 15 | Perfusate inputs / outputs | 1 Glc Tracer IN | Glc.tracer -> Glc.perf |
| 16 | Perfusate inputs / outputs | 1 NA Glc IN | Glc.source -> Glc.perf |
| 17 | Perfusate inputs / outputs | 1 Total Glc IN | Glc.perf -> Glc.aff |
| 18 | Perfusate inputs / outputs | 1 Glc balance net | Glc <-> Glc.aff |
| 19 | Perfusate inputs / outputs | 1 Glc balance exch | Glc <-> Glc.aff |
| 20 | Perfusate inputs / outputs | 1 Glc effluent | Glc.aff -> Glc.eff |
| 21 | Perfusate inputs / outputs | 1 Total Glc OUT | Glc.eff -> Glc.sink |
| 22 | Perfusate inputs / outputs | 1 Glycerol source | Glycerol.source -> Glycerol.aff |
| 23 | Perfusate inputs / outputs | Glycerol.eff | Glycerol.aff -> Glycerol.eff |
| 24 | Perfusate inputs / outputs | Glycerol.sink | Glycerol.eff -> Glycerol.sink |
| 25 | Perfusate inputs / outputs | Bicarb source | Bicarb.source -> CO2.aff |
| 26 | Perfusate inputs / outputs | Bicarb Sink | CO2.aff -> Bicarb.sink |
| 27 | β-oxidation / acetyl-CoA | A1 | Fat -> AcCoA |
| 28 | TCA cycle / anaplerosis / $CO_2$ | A2 | Pyr -> AcCoA + CO2 |
| 29 | TCA cycle / anaplerosis / $CO_2$ | A3 | Oac + AcCoA -> Cit |
| 30 | TCA cycle / anaplerosis / $CO_2$ | A4 | Pyr + CO2 -> Oac |
| 31 | TCA cycle / anaplerosis / $CO_2$ | A5 net | Cit <-> Akg + CO2 |
| 32 | TCA cycle / anaplerosis / $CO_2$ | A5 exch | Cit <-> Akg + CO2 |
| 33 | TCA cycle / anaplerosis / $CO_2$ | A6 | Akg -> SucCoA + CO2 |
| 34 | TCA cycle / anaplerosis / $CO_2$ | A7 net | Glu <-> Akg |
| 35 | TCA cycle / anaplerosis / $CO_2$ | A7 exch | Glu <-> Akg |
| 36 | TCA cycle / anaplerosis / $CO_2$ | A8 | SucCoA -> Suc |
| 37 | TCA cycle / anaplerosis / $CO_2$ | A9 net | Suc <-> Fum |
| 38 | TCA cycle / anaplerosis / $CO_2$ | A9 exch | Suc <-> Fum |
| 39 | TCA cycle / anaplerosis / $CO_2$ | A10 net | Fum <-> Mal |
| 40 | TCA cycle / anaplerosis / $CO_2$ | A10 exch | Fum <-> Mal |
| 41 | TCA cycle / anaplerosis / $CO_2$ | A11 net | Mal <-> Oac |
| 42 | TCA cycle / anaplerosis / $CO_2$ | A11 exch | Mal <-> Oac |
| 43 | TCA cycle / anaplerosis / $CO_2$ | A12 net | Oac <-> Asp |
| 44 | TCA cycle / anaplerosis / $CO_2$ | A12 exch | Oac <-> Asp |
| 45 | TCA cycle / anaplerosis / $CO_2$ | T5 net | CO2.aff <-> CO2 |
| 46 | TCA cycle / anaplerosis / $CO_2$ | T5 exch | CO2.aff <-> CO2 |
| 47 | Pyruvate / lactate / alanine exchange | B17 net | Ala <-> Pyr |
| 48 | Pyruvate / lactate / alanine exchange | B17 exch | Ala <-> Pyr |
| 49 | Pyruvate / lactate / alanine exchange | B18 net | Lac <-> Pyr |
| 50 | Pyruvate / lactate / alanine exchange | B18 exch | Lac <-> Pyr |

| Order | Pathway category | Reaction label | Reaction equation |
|---|---|---|---|
| 51 | Gluconeogenesis / glycolysis / glycogen | A27* | Oac -> PEP + CO2 |
| 52 | Gluconeogenesis / glycolysis / glycogen | B28 | PEP -> Pyr |
| 53 | Gluconeogenesis / glycolysis / glycogen | B29 net | PEP <-> BPG |
| 54 | Gluconeogenesis / glycolysis / glycogen | B29 exch | PEP <-> BPG |
| 55 | Gluconeogenesis / glycolysis / glycogen | B30 net | BPG <-> GAP |
| 56 | Gluconeogenesis / glycolysis / glycogen | B30 exch | BPG <-> GAP |
| 57 | Gluconeogenesis / glycolysis / glycogen | B31 | Glycerol.aff -> G3P |
| 58 | Gluconeogenesis / glycolysis / glycogen | B32 | G3P -> DHAP |
| 59 | Gluconeogenesis / glycolysis / glycogen | B33 net | DHAP <-> GAP |
| 60 | Gluconeogenesis / glycolysis / glycogen | B33 exch | DHAP <-> GAP |
| 61 | Gluconeogenesis / glycolysis / glycogen | B34 net | DHAP + GAP <-> G6P |
| 62 | Gluconeogenesis / glycolysis / glycogen | B34 exch | DHAP + GAP <-> G6P |
| 63 | Gluconeogenesis / glycolysis / glycogen | B35 | G6P -> Glc |
| 64 | Gluconeogenesis / glycolysis / glycogen | B36 | Glycogen -> G6P |
| 65 | Measurement / contribution bookkeeping | Contribution Ala | 0*Ala -> Ala.s |
| 66 | Measurement / contribution bookkeeping | Contribution Ala tot | Ala.s -> Ala.sink |
| 67 | Measurement / contribution bookkeeping | Contributino Akg | 0*Akg -> Akg.s |
| 68 | Measurement / contribution bookkeeping | Contribution Akg total | Akg.s -> Akg.sink |
| 69 | Measurement / contribution bookkeeping | Contribution Asp | 0*Asp -> Asp.s |
| 70 | Measurement / contribution bookkeeping | Contribution Asp total | Asp.s -> Asp.sink |
| 71 | Measurement / contribution bookkeeping | Contribution Cit | 0*Cit -> Cit.s |
| 72 | Measurement / contribution bookkeeping | Contribution Cit Tot | Cit.s -> Cit.sink |
| 73 | Measurement / contribution bookkeeping | Contribution Fum sourc | 0*Fum -> Fum.s |
| 74 | Measurement / contribution bookkeeping | Contribution Fum total | Fum.s -> Fum.sink |
| 75 | Measurement / contribution bookkeeping | Contribution Glu | 0*Glu -> Glu.s |
| 76 | Measurement / contribution bookkeeping | Contribution Glu tot | Glu.s -> Glu.sink |
| 77 | Measurement / contribution bookkeeping | Contribution Lac | 0*Lac -> Lac.s |
| 78 | Measurement / contribution bookkeeping | Contribution Lac.c | 0*Lac.c -> Lac.s |
| 79 | Measurement / contribution bookkeeping | Constribution Lac total | Lac.s -> Lac.sink |
| 80 | Measurement / contribution bookkeeping | Contribution Mal | 0*Mal -> Mal.s |

| Order | Pathway category | Reaction label | Reaction equation |
|---|---|---|---|
| 81 | Measurement / contribution bookkeeping | Contribution Mal tot | Mal.s -> Mal.sink |
| 82 | Measurement / contribution bookkeeping | Contribution Pyr | 0*Pyr -> Pyr.s |
| 83 | Measurement / contribution bookkeeping | Contribution Pyr total | Pyr.s -> Pyr.sink |
| 84 | Measurement / contribution bookkeeping | Contribution Suc source | 0*Suc -> Suc.s |
| 85 | Measurement / contribution bookkeeping | Contribution suc tot | Suc.s -> Suc.sink |
| 86 | Measurement / contribution bookkeeping | Z13 net | Asp.inert <-> Asp + Inertasp |
| 87 | Measurement / contribution bookkeeping | Z13 exch | Asp.inert <-> Asp + Inertasp |
| 88 | Measurement / contribution bookkeeping | Z14 net | Glu.inert <-> Glu + InertGlu |
| 89 | Measurement / contribution bookkeeping | Z14 exch | Glu.inert <-> Glu + InertGlu |
| 90 | Measurement / contribution bookkeeping | Z1 | 0*Ala.cold -> Ala.s |
| 91 | Measurement / contribution bookkeeping | Z2 | 0*Asp.cold -> Asp.s |

#### S1C. Two-compartment model

| Order | Pathway category | Reaction label | Reaction equation |
|---|---|---|---|
| 1 | Perfusate inputs / outputs | 1 Lac tracer IN | Lac.tracer -> Lac.perf + 0.1*PyrTr |
| 2 | Perfusate inputs / outputs | 1 NA Lac IN | Lac.source -> Lac.perf + 0.1*PyrNA |
| 3 | Perfusate inputs / outputs | 1 Total Lac IN | Lac.perf -> Lac.aff |
| 4 | Perfusate inputs / outputs | 1 Lac Balance net | Lac.aff <-> Lac |
| 5 | Perfusate inputs / outputs | 1 Lac Balance exch | Lac.aff <-> Lac |
| 6 | Perfusate inputs / outputs | 1 Lac effluent | Lac.aff -> Lac.eff |
| 7 | Perfusate inputs / outputs | 1 Total Lac OUT | Lac.eff -> Lac.sink |
| 8 | Perfusate inputs / outputs | 1 Pyr tracer IN | Pyr.tracer + PyrTr -> Pyr.perf |
| 9 | Perfusate inputs / outputs | 1 NA Pyr IN | Pyr.source + PyrNA -> Pyr.perf |
| 10 | Perfusate inputs / outputs | 1 Total Pyr IN | Pyr.perf -> Pyr.aff |
| 11 | Perfusate inputs / outputs | 1 Pyr Balance net | Pyr.aff <-> Pyr |
| 12 | Perfusate inputs / outputs | 1 Pyr Balance exch | Pyr.aff <-> Pyr |
| 13 | Perfusate inputs / outputs | 1 Pyr effluent | Pyr.aff -> Pyr.eff |
| 14 | Perfusate inputs / outputs | 1 Total Pyr OUT | Pyr.eff -> Pyr.sink |
| 15 | Perfusate inputs / outputs | 1 Glc Tracer IN | Glc.tracer -> Glc.perf |
| 16 | Perfusate inputs / outputs | 1 NA Glc IN | Glc.source -> Glc.perf |
| 17 | Perfusate inputs / outputs | 1 Total Glc IN | Glc.perf -> Glc.aff |
| 18 | Perfusate inputs / outputs | 1 Glc balance net | Glc <-> Glc.aff |
| 19 | Perfusate inputs / outputs | 1 Glc balance exch | Glc <-> Glc.aff |
| 20 | Perfusate inputs / outputs | 1 Glc effluent | Glc.aff -> Glc.eff |
| 21 | Perfusate inputs / outputs | 1 Total Glc OUT | Glc.eff -> Glc.sink |
| 22 | Perfusate inputs / outputs | 1 Glycerol source | Glycerol.source -> Glycerol.aff |
| 23 | Perfusate inputs / outputs | Glycerol.eff | Glycerol.aff -> Glycerol.eff |
| 24 | Perfusate inputs / outputs | Glycerol.sink | Glycerol.eff -> Glycerol.sink |
| 25 | Perfusate inputs / outputs | Bicarb source | Bicarb.source -> CO2.aff |
| 26 | Perfusate inputs / outputs | Bicarb Sink | CO2.aff -> Bicarb.sink |
| 27 | β-oxidation / acetyl-CoA | A1 | Fat -> AcCoA |
| 28 | TCA cycle / anaplerosis / $CO_2$ | A2 | Pyr -> AcCoA + CO2 |
| 29 | TCA cycle / anaplerosis / $CO_2$ | A3 | Oac + AcCoA -> Cit |

| Order | Pathway category | Reaction label | Reaction equation |
|---|---|---|---|
| 30 | TCA cycle / anaplerosis / $CO_2$ | A4 | Pyr + CO2 -> Oac |
| 31 | TCA cycle / anaplerosis / $CO_2$ | A5 net | Cit <-> Akg + CO2 |
| 32 | TCA cycle / anaplerosis / $CO_2$ | A5 exch | Cit <-> Akg + CO2 |
| 33 | TCA cycle / anaplerosis / $CO_2$ | A6 | Akg -> SucCoA + CO2 |
| 34 | TCA cycle / anaplerosis / $CO_2$ | A7 net | Glu <-> Akg |
| 35 | TCA cycle / anaplerosis / $CO_2$ | A7 exch | Glu <-> Akg |
| 36 | TCA cycle / anaplerosis / $CO_2$ | A8 | SucCoA -> Suc |
| 37 | TCA cycle / anaplerosis / $CO_2$ | A9 net | Suc <-> Fum |
| 38 | TCA cycle / anaplerosis / $CO_2$ | A9 exch | Suc <-> Fum |
| 39 | TCA cycle / anaplerosis / $CO_2$ | A10 net | Fum <-> Mal |
| 40 | TCA cycle / anaplerosis / $CO_2$ | A10 exch | Fum <-> Mal |
| 41 | TCA cycle / anaplerosis / $CO_2$ | A11 net | Mal <-> Oac |
| 42 | TCA cycle / anaplerosis / $CO_2$ | A11 exch | Mal <-> Oac |
| 43 | TCA cycle / anaplerosis / $CO_2$ | A12 net | Oac <-> Asp |
| 44 | TCA cycle / anaplerosis / $CO_2$ | A12 exch | Oac <-> Asp |
| 45 | TCA cycle / anaplerosis / $CO_2$ | T5 net | CO2.aff <-> CO2 |
| 46 | TCA cycle / anaplerosis / $CO_2$ | T5 exch | CO2.aff <-> CO2 |
| 47 | Pyruvate / lactate / alanine exchange | B17 net | Ala <-> Pyr |
| 48 | Pyruvate / lactate / alanine exchange | B17 exch | Ala <-> Pyr |
| 49 | Pyruvate / lactate / alanine exchange | B18 net | Lac <-> Pyr |
| 50 | Pyruvate / lactate / alanine exchange | B18 exch | Lac <-> Pyr |
| 51 | Gluconeogenesis / glycolysis / glycogen | A27* | Oac -> PEP + CO2 |
| 52 | Gluconeogenesis / glycolysis / glycogen | B28 | PEP -> Pyr |
| 53 | Gluconeogenesis / glycolysis / glycogen | B29 net | PEP <-> BPG |
| 54 | Gluconeogenesis / glycolysis / glycogen | B29 exch | PEP <-> BPG |
| 55 | Gluconeogenesis / glycolysis / glycogen | B30 net | BPG <-> GAP |
| 56 | Gluconeogenesis / glycolysis / glycogen | B30 exch | BPG <-> GAP |
| 57 | Gluconeogenesis / glycolysis / glycogen | B31 | Glycerol.aff -> G3P |
| 58 | Gluconeogenesis / glycolysis / glycogen | B32 | G3P -> DHAP |
| 59 | Gluconeogenesis / glycolysis / glycogen | B33 net | DHAP <-> GAP |
| 60 | Gluconeogenesis / glycolysis / glycogen | B33 exch | DHAP <-> GAP |
| 61 | Gluconeogenesis / glycolysis / glycogen | B34 net | DHAP + GAP <-> G6P |
| 62 | Gluconeogenesis / glycolysis / glycogen | B34 exch | DHAP + GAP <-> G6P |
| 63 | Gluconeogenesis / glycolysis / glycogen | B35 net | G6P <-> Glc |
| 64 | Gluconeogenesis / glycolysis / glycogen | B35 exch | G6P <-> Glc |
| 65 | Gluconeogenesis / glycolysis / glycogen | B36 | Glycogen -> G6P |
| 66 | Secondary compartment | C1 | Glc.aff -> Glc.c |
| 67 | Secondary compartment | C3 | Glc.c -> G6P.c |
| 68 | Secondary compartment | C4 net | G6P.c <-> DHAP.c + GAP.c |
| 69 | Secondary compartment | C4 exch | G6P.c <-> DHAP.c + GAP.c |
| 70 | Secondary compartment | C5 net | DHAP.c <-> GAP.c |

| Order | Pathway category | Reaction label | Reaction equation |
| --- | --- | --- | --- |
| 71 | Secondary compartment | C5 exch | DHAP.c <-> GAP.c |
| 72 | Secondary compartment | C6 | GAP.c -> BPG.c |
| 73 | Secondary compartment | C7 | BPG.c -> PEP.c |
| 74 | Secondary compartment | C8 | PEP.c -> Pyr.c |
| 75 | Secondary compartment | C9 net | Lac.c <-> Pyr.c |
| 76 | Secondary compartment | C9 exch | Lac.c <-> Pyr.c |
| 77 | Secondary compartment | C10 net | Ala.c <-> Pyr.c |
| 78 | Secondary compartment | C10 exch | Ala.c <-> Pyr.c |
| 79 | Secondary compartment | C11 | Lac.aff -> Lac.c |
| 80 | Secondary compartment | C12 | Lac.c -> Lac.eff |
| 81 | Secondary compartment | C13 | Pyr.aff -> Pyr.c |
| 82 | Secondary compartment | C14 | Pyr.c -> Pyr.eff |
| 83 | Secondary compartment | C15 | Pyr.c -> AcCoA.c + CO2.c |
| 84 | Secondary compartment | C18 net | Cit.c <-> Akg.c + CO2.c |
| 85 | Secondary compartment | C18 exch | Cit.c <-> Akg.c + CO2.c |
| 86 | Secondary compartment | C19 net | Glu.c <-> Akg.c |
| 87 | Secondary compartment | C19 exch | Glu.c <-> Akg.c |
| 88 | Secondary compartment | C20 | Oac.c + AcCoA.c -> Cit.c |
| 89 | Secondary compartment | C21 | Akg.c -> SucCoA.c + CO2.c |
| 90 | Secondary compartment | C22 | SucCoA.c -> Suc.c |
| 91 | Secondary compartment | C23 net | Suc.c <-> Fum.c |
| 92 | Secondary compartment | C23 exch | Suc.c <-> Fum.c |
| 93 | Secondary compartment | C24 net | Fum.c <-> Mal.c |
| 94 | Secondary compartment | C24 exch | Fum.c <-> Mal.c |
| 95 | Secondary compartment | C25 net | Mal.c <-> Oac.c |
| 96 | Secondary compartment | C25 exch | Mal.c <-> Oac.c |
| 97 | Secondary compartment | C26 net | Oac.c <-> Asp.c |
| 98 | Secondary compartment | C26 exch | Oac.c <-> Asp.c |
| 99 | Secondary compartment | C28 net | Pyr.c + CO2.c <-> Mal.c + ME.c |
| 100 | Secondary compartment | C28 exch | Pyr.c + CO2.c <-> Mal.c + ME.c |
| 101 | Secondary compartment | C29 | Mal.c -> Pyr.c + CO2.c |
| 102 | Secondary compartment | C34 net | CO2.aff <-> CO2.c |
| 103 | Secondary compartment | C34 exch | CO2.aff <-> CO2.c |
| 104 | Measurement / contribution bookkeeping | Contribution Ala | 0*Ala -> Ala.s |
| 105 | Measurement / contribution bookkeeping | Contribution Ala.c | 0*Ala.c -> Ala.s |
| 106 | Measurement / contribution bookkeeping | Contribution Ala tot | Ala.s -> Ala.sink |
| 107 | Measurement / contribution bookkeeping | Contributino Akg | 0*Akg -> Akg.s |
| 108 | Measurement / contribution bookkeeping | Contribution Akg.c | 0*Akg.c -> Akg.s |
| 109 | Measurement / contribution bookkeeping | Contribution Akg total | Akg.s -> Akg.sink |
| 110 | Measurement / contribution bookkeeping | Contribution Asp | 0*Asp -> Asp.s |
| 111 | Measurement / contribution bookkeeping | Contribution Asp.c | 0*Asp.c -> Asp.s |
| 112 | Measurement / contribution bookkeeping | Contribution Asp total | Asp.s -> Asp.sink |
| 113 | Measurement / contribution bookkeeping | Contribution Cit | 0*Cit -> Cit.s |
| 114 | Measurement / contribution bookkeeping | Contribution Cit.c | 0*Cit.c -> Cit.s |

| Order | Pathway category | Reaction label | Reaction equation |
|---|---|---|---|
| 115 | Measurement / contribution bookkeeping | Contribution Cit Tot | Cit.s -> Cit.sink |
| 116 | Measurement / contribution bookkeeping | Contribution Fum sourc | 0*Fum -> Fum.s |
| 117 | Measurement / contribution bookkeeping | Contribution Fum.c | 0*Fum.c -> Fum.s |
| 118 | Measurement / contribution bookkeeping | Contribution Fum total | Fum.s -> Fum.sink |
| 119 | Measurement / contribution bookkeeping | Contribution Glu | 0*Glu -> Glu.s |
| 120 | Measurement / contribution bookkeeping | Contribution Glu.c | 0*Glu.c -> Glu.s |
| 121 | Measurement / contribution bookkeeping | Contribution Glu tot | Glu.s -> Glu.sink |
| 122 | Measurement / contribution bookkeeping | Contribution Lac | 0*Lac -> Lac.s |
| 123 | Measurement / contribution bookkeeping | Contribution Lac.c | 0*Lac.c -> Lac.s |
| 124 | Measurement / contribution bookkeeping | Constribution Lac total | Lac.s -> Lac.sink |
| 125 | Measurement / contribution bookkeeping | Contribution Mal | 0*Mal -> Mal.s |
| 126 | Measurement / contribution bookkeeping | Contribution Mal.c | 0*Mal.c -> Mal.s |
| 127 | Measurement / contribution bookkeeping | Contribution Mal tot | Mal.s -> Mal.sink |
| 128 | Measurement / contribution bookkeeping | Contribution Pyr | 0*Pyr -> Pyr.s |
| 129 | Measurement / contribution bookkeeping | Contribution Pyr total | Pyr.s -> Pyr.sink |
| 130 | Measurement / contribution bookkeeping | Contribution Suc source | 0*Suc -> Suc.s |
| 131 | Measurement / contribution bookkeeping | Contribution Suc.c | 0*Suc.c -> Suc.s |
| 132 | Measurement / contribution bookkeeping | Contribution suc tot | Suc.s -> Suc.sink |
| 133 | Measurement / contribution bookkeeping | Z1 | 0*Ala.cold -> Ala.s |
| 134 | Measurement / contribution bookkeeping | Z2 | 0*Asp.cold -> Asp.s |
| 135 | Measurement / contribution bookkeeping | Z3 | 0*Glu.cold -> Glu.s |
| 136 | Measurement / contribution bookkeeping | Z5 | Aspc.sourcec -> Asp.c |
| 137 | Measurement / contribution bookkeeping | Z6 | Gluc.sourcec -> Glu.c |

#### Table S2. Measurement and boundary-flux constraints

*“Included” indicates whether each measurement or boundary flux was used as a fitting constraint in that model.*

##### S2A. Glucose fragment model

| Order | Measurement / boundary constraint | Constraint type | Included |
|---|---|---|---|
| 1 | Liver Pyr174 | Intracellular MID | No |
| 2 | Liver Lac233 | Intracellular MID | No |
| 3 | Liver Lac261 | Intracellular MID | No |
| 4 | Liver Ala232 | Intracellular MID | No |
| 5 | Liver Ala260 | Intracellular MID | No |
| 6 | Liver Urea231 | Intracellular MID | No |
| 7 | Liver Suc289 | Intracellular MID | No |
| 8 | Liver Fum287 | Intracellular MID | No |
| 9 | Liver OAA332 | Intracellular MID | No |
| 10 | Liver Akg346 | Intracellular MID | No |
| 11 | Liver Malate391 | Intracellular MID | No |
| 12 | Liver Malate419 | Intracellular MID | No |
| 13 | Liver Asp302 | Intracellular MID | No |
| 14 | Liver Asp390 | Intracellular MID | No |
| 15 | Liver Asp418 | Intracellular MID | No |
| 16 | Liver PEP453 | Intracellular MID | No |
| 17 | Liver DHAP484 | Intracellular MID | No |
| 18 | Liver G3P571 | Intracellular MID | No |
| 19 | Liver Cit431 | Intracellular MID | No |
| 20 | Liver Cit459 | Intracellular MID | No |
| 21 | Perfusate Glc (DIO301) | Perfusate MID | No |
| 22 | Perfusate Glc (MOX145) | Perfusate MID | No |
| 23 | Perfusate Glc (Aldo173) | Perfusate MID | No |
| 24 | Perfusate Glc (Aldo259) | Perfusate MID | No |
| 25 | Perfusate Glc (Aldo284) | Perfusate MID | No |
| 26 | Perfusate Glc (Aldo370) | Perfusate MID | No |
| 27 | Effluent Glc (DIO301) | Effluent MID | Yes |
| 28 | Effluent Glc (MOX145) | Effluent MID | Yes |
| 29 | Effluent Glc (Aldo173) | Effluent MID | Yes |
| 30 | Effluent Glc (Aldo259) | Effluent MID | Yes |
| 31 | Effluent Glc (Aldo284) | Effluent MID | Yes |
| 32 | Effluent Glc (Aldo370) | Effluent MID | Yes |
| 33 | Perfusate Pyr174 | Perfusate MID | No |
| 34 | Perfusate Lac233 | Perfusate MID | No |
| 35 | Perfusate Lac261 | Perfusate MID | No |
| 36 | Effluent Pyr174 | Effluent MID | No |
| 37 | Effluent Lac233 | Effluent MID | No |
| 38 | Effluent Lac261 | Effluent MID | No |
| 39 | Liver Glu330 | Intracellular MID | No |
| 40 | Liver Glu432 | Intracellular MID | No |
| 41 | 1 Total Glc IN | Measured flux / boundary constraint | Yes |
| 42 | 1 Total Glc OUT | Measured flux / boundary constraint | Yes |
| 43 | 1 Total Lac IN | Measured flux / boundary constraint | No |
| 44 | 1 Total Lac OUT | Measured flux / boundary constraint | No |
| 45 | 1 Total Pyr IN | Measured flux / boundary constraint | No |
| 46 | 1 Total Pyr Out | Measured flux / boundary constraint | No |
| 47 | 1 Glycerol Source | Measured flux / boundary constraint | Yes |

| Order | Measurement / boundary constraint | Constraint type | Included |
|---|---|---|---|
| 48 | Glycerol.eff | Measured flux / boundary constraint | Yes |
| 49 | 1 NA Lac IN | Measured flux / boundary constraint | Yes |
| 50 | 1 Total Lac IN | Measured flux / boundary constraint | Yes |
| 51 | 1 NA Pyr IN | Measured flux / boundary constraint | Yes |

#### S2B. One-compartment model, full constraint set

| Order | Measurement / boundary constraint | Constraint type | Included |
|---|---|---|---|
| 1 | Liver Pyr174 | Intracellular MID | No |
| 2 | Liver Lac233 | Intracellular MID | Yes |
| 3 | Liver Lac261 | Intracellular MID | Yes |
| 4 | Liver Ala232 | Intracellular MID | Yes |
| 5 | Liver Ala260 | Intracellular MID | Yes |
| 6 | Liver Urea231 | Intracellular MID | No |
| 7 | Liver Suc289 | Intracellular MID | Yes |
| 8 | Liver Fum287 | Intracellular MID | Yes |
| 9 | Liver OAA332 | Intracellular MID | No |
| 10 | Liver Akg346 | Intracellular MID | Yes |
| 11 | Liver Malate391 | Intracellular MID | Yes |
| 12 | Liver Malate419 | Intracellular MID | Yes |
| 13 | Liver Asp302 | Intracellular MID | Yes |
| 14 | Liver Asp390 | Intracellular MID | Yes |
| 15 | Liver Asp418 | Intracellular MID | Yes |
| 16 | Liver PEP453 | Intracellular MID | No |
| 17 | Liver DHAP484 | Intracellular MID | No |
| 18 | Liver G3P571 | Intracellular MID | No |
| 19 | Liver Cit431 | Intracellular MID | No |
| 20 | Liver Cit459 | Intracellular MID | Yes |
| 21 | Perfusate Glc (DIO301) | Perfusate MID | No |
| 22 | Perfusate Glc (MOX145) | Perfusate MID | No |
| 23 | Perfusate Glc (Aldo173) | Perfusate MID | No |
| 24 | Perfusate Glc (Aldo259) | Perfusate MID | No |
| 25 | Perfusate Glc (Aldo284) | Perfusate MID | No |
| 26 | Perfusate Glc (Aldo370) | Perfusate MID | No |
| 27 | Effluent Glc (DIO301) | Effluent MID | Yes |
| 28 | Effluent Glc (MOX145) | Effluent MID | Yes |
| 29 | Effluent Glc (Aldo173) | Effluent MID | Yes |
| 30 | Effluent Glc (Aldo259) | Effluent MID | Yes |
| 31 | Effluent Glc (Aldo284) | Effluent MID | Yes |
| 32 | Effluent Glc (Aldo370) | Effluent MID | Yes |
| 33 | Perfusate Pyr174 | Perfusate MID | No |
| 34 | Perfusate Lac233 | Perfusate MID | Yes |
| 35 | Perfusate Lac261 | Perfusate MID | Yes |
| 36 | Effluent Pyr174 | Effluent MID | Yes |
| 37 | Effluent Lac233 | Effluent MID | Yes |
| 38 | Effluent Lac261 | Effluent MID | Yes |
| 39 | Liver Glu330 | Intracellular MID | Yes |
| 40 | Liver Glu432 | Intracellular MID | Yes |
| 41 | 1 Total Glc IN | Measured flux / boundary constraint | Yes |
| 42 | 1 Total Glc OUT | Measured flux / boundary constraint | Yes |
| 43 | 1 Total Lac IN | Measured flux / boundary constraint | No |
| 44 | 1 Total Lac OUT | Measured flux / boundary constraint | No |

| Order | Measurement / boundary constraint | Constraint type | Included |
|---|---|---|---|
| 45 | 1 Total Pyr IN | Measured flux / boundary constraint | No |
| 46 | 1 Total Pyr Out | Measured flux / boundary constraint | No |
| 47 | 1 Glycerol Source | Measured flux / boundary constraint | Yes |
| 48 | Glycerol.eff | Measured flux / boundary constraint | Yes |
| 49 | 1 NA Lac IN | Measured flux / boundary constraint | Yes |
| 50 | 1 Total Lac IN | Measured flux / boundary constraint | Yes |
| 51 | 1 NA Pyr IN | Measured flux / boundary constraint | Yes |

#### S2C. One-compartment model, reduced constraint set

| Order | Measurement / boundary constraint | Constraint type | Included |
|---|---|---|---|
| 1 | Liver Pyr174 | Intracellular MID | No |
| 2 | Liver Lac233 | Intracellular MID | Yes |
| 3 | Liver Lac261 | Intracellular MID | Yes |
| 4 | Liver Ala232 | Intracellular MID | Yes |
| 5 | Liver Ala260 | Intracellular MID | Yes |
| 6 | Liver Urea231 | Intracellular MID | No |
| 7 | Liver Suc289 | Intracellular MID | No |
| 8 | Liver Fum287 | Intracellular MID | No |
| 9 | Liver OAA332 | Intracellular MID | No |
| 10 | Liver Akg346 | Intracellular MID | Yes |
| 11 | Liver Malate391 | Intracellular MID | No |
| 12 | Liver Malate419 | Intracellular MID | No |
| 13 | Liver Asp302 | Intracellular MID | No |
| 14 | Liver Asp390 | Intracellular MID | No |
| 15 | Liver Asp418 | Intracellular MID | No |
| 16 | Liver PEP453 | Intracellular MID | No |
| 17 | Liver DHAP484 | Intracellular MID | No |
| 18 | Liver G3P571 | Intracellular MID | No |
| 19 | Liver Cit431 | Intracellular MID | No |
| 20 | Liver Cit459 | Intracellular MID | Yes |
| 21 | Perfusate Glc (DIO301) | Perfusate MID | No |
| 22 | Perfusate Glc (MOX145) | Perfusate MID | No |
| 23 | Perfusate Glc (Aldo173) | Perfusate MID | No |
| 24 | Perfusate Glc (Aldo259) | Perfusate MID | No |
| 25 | Perfusate Glc (Aldo284) | Perfusate MID | No |
| 26 | Perfusate Glc (Aldo370) | Perfusate MID | No |
| 27 | Effluent Glc (DIO301) | Effluent MID | Yes |
| 28 | Effluent Glc (MOX145) | Effluent MID | Yes |
| 29 | Effluent Glc (Aldo173) | Effluent MID | Yes |
| 30 | Effluent Glc (Aldo259) | Effluent MID | Yes |
| 31 | Effluent Glc (Aldo284) | Effluent MID | Yes |
| 32 | Effluent Glc (Aldo370) | Effluent MID | Yes |
| 33 | Perfusate Pyr174 | Perfusate MID | No |
| 34 | Perfusate Lac233 | Perfusate MID | Yes |
| 35 | Perfusate Lac261 | Perfusate MID | Yes |
| 36 | Effluent Pyr174 | Effluent MID | Yes |
| 37 | Effluent Lac233 | Effluent MID | Yes |
| 38 | Effluent Lac261 | Effluent MID | Yes |
| 39 | Liver Glu330 | Intracellular MID | Yes |
| 40 | Liver Glu432 | Intracellular MID | Yes |
| 41 | 1 Total Glc IN | Measured flux / boundary constraint | Yes |

| Order | Measurement / boundary constraint | Constraint type | Included |
|---|---|---|---|
| 42 | 1 Total Glc OUT | Measured flux / boundary constraint | Yes |
| 43 | 1 Total Lac IN | Measured flux / boundary constraint | No |
| 44 | 1 Total Lac OUT | Measured flux / boundary constraint | No |
| 45 | 1 Total Pyr IN | Measured flux / boundary constraint | No |
| 46 | 1 Total Pyr Out | Measured flux / boundary constraint | No |
| 47 | 1 Glycerol Source | Measured flux / boundary constraint | Yes |
| 48 | Glycerol.eff | Measured flux / boundary constraint | Yes |
| 49 | 1 NA Lac IN | Measured flux / boundary constraint | Yes |
| 50 | 1 Total Lac IN | Measured flux / boundary constraint | Yes |
| 51 | 1 NA Pyr IN | Measured flux / boundary constraint | Yes |

#### S2D. Two-compartment model

| Order | Measurement / boundary constraint | Constraint type | Included |
|---|---|---|---|
| 1 | Liver Pyr174 | Intracellular MID | No |
| 2 | Liver Lac233 | Intracellular MID | Yes |
| 3 | Liver Lac261 | Intracellular MID | Yes |
| 4 | Liver Ala232 | Intracellular MID | Yes |
| 5 | Liver Ala260 | Intracellular MID | Yes |
| 6 | Liver Urea231 | Intracellular MID | No |
| 7 | Liver Suc289 | Intracellular MID | Yes |
| 8 | Liver Fum287 | Intracellular MID | Yes |
| 9 | Liver OAA332 | Intracellular MID | No |
| 10 | Liver Akg346 | Intracellular MID | Yes |
| 11 | Liver Malate391 | Intracellular MID | Yes |
| 12 | Liver Malate419 | Intracellular MID | Yes |
| 13 | Liver Asp302 | Intracellular MID | Yes |
| 14 | Liver Asp390 | Intracellular MID | Yes |
| 15 | Liver Asp418 | Intracellular MID | Yes |
| 16 | Liver PEP453 | Intracellular MID | No |
| 17 | Liver DHAP484 | Intracellular MID | No |
| 18 | Liver G3P571 | Intracellular MID | No |
| 19 | Liver Cit431 | Intracellular MID | No |
| 20 | Liver Cit459 | Intracellular MID | Yes |
| 21 | Perfusate Glc (DIO301) | Perfusate MID | No |
| 22 | Perfusate Glc (MOX145) | Perfusate MID | No |
| 23 | Perfusate Glc (Aldo173) | Perfusate MID | No |
| 24 | Perfusate Glc (Aldo259) | Perfusate MID | No |
| 25 | Perfusate Glc (Aldo284) | Perfusate MID | No |
| 26 | Perfusate Glc (Aldo370) | Perfusate MID | No |
| 27 | Effluent Glc (DIO301) | Effluent MID | Yes |
| 28 | Effluent Glc (MOX145) | Effluent MID | Yes |
| 29 | Effluent Glc (Aldo173) | Effluent MID | Yes |
| 30 | Effluent Glc (Aldo259) | Effluent MID | Yes |
| 31 | Effluent Glc (Aldo284) | Effluent MID | Yes |
| 32 | Effluent Glc (Aldo370) | Effluent MID | Yes |
| 33 | Perfusate Pyr174 | Perfusate MID | No |
| 34 | Perfusate Lac233 | Perfusate MID | Yes |
| 35 | Perfusate Lac261 | Perfusate MID | Yes |
| 36 | Effluent Pyr174 | Effluent MID | Yes |
| 37 | Effluent Lac233 | Effluent MID | Yes |
| 38 | Effluent Lac261 | Effluent MID | Yes |

| Order | Measurement / boundary constraint | Constraint type | Included |
|---|---|---|---|
| 39 | Liver Glu330 | Intracellular MID | Yes |
| 40 | Liver Glu432 | Intracellular MID | Yes |
| 41 | 1 Total Glc IN | Measured flux / boundary constraint | Yes |
| 42 | 1 Total Glc OUT | Measured flux / boundary constraint | Yes |
| 43 | 1 Total Lac IN | Measured flux / boundary constraint | No |
| 44 | 1 Total Lac OUT | Measured flux / boundary constraint | No |
| 45 | 1 Total Pyr IN | Measured flux / boundary constraint | No |
| 46 | 1 Total Pyr Out | Measured flux / boundary constraint | No |
| 47 | 1 Glycerol Source | Measured flux / boundary constraint | Yes |
| 48 | Glycerol.eff | Measured flux / boundary constraint | Yes |
| 49 | 1 NA Lac IN | Measured flux / boundary constraint | Yes |
| 50 | 1 Total Lac IN | Measured flux / boundary constraint | Yes |

#### S2E. Measurement, derivative and fragment definitions

*Maximum isotopologue is set by the number of labeled carbon positions in each fragment (for example, three labeled carbons permit M+0 to M+3). "Active" indicates use as a constraint in the two-compartment model.*

| INCA measurement ID | Pool ID | Derivative / fragment | m/z | Labeled Atom IDs | Max labeled C | Max isotopologue | Unlabeled atoms | Active |
|---|---|---|---|---|---|---|---|---|
| Liver Pyr174 | Pyr.s | Pyr174 | 174 | 1 2 3 | 3 | M+3 | C3H12O3NSi | No |
| Liver Lac233 | Lac.s | Lac233 | 233 | 2 3 | 2 | M+2 | C8H25O2Si2 | Yes |
| Liver Lac261 | Lac.s | Lac261 | 261 | 1 2 3 | 3 | M+3 | C8H25O3Si2 | Yes |
| Liver Ala232 | Ala.s | Ala232 | 232 | 2 3 | 2 | M+2 | C8H26ONSi2 | Yes |
| Liver Ala260 | Ala.s | Ala260 | 260 | 1 2 3 | 3 | M+3 | C8H26O2NSi2 | Yes |
| Liver Urea231 | CO2 | Urea231 | 231 | 1 | 1 | M+1 | C8H23N2OSi2 | No |
| Liver Suc289 | Suc.s | Suc289 | 289 | 1 2 3 4 | 4 | M+4 | C8H25O4Si2 | Yes |
| Liver Fum287 | Fum.s | Fum287 | 287 | 1 2 3 4 | 4 | M+4 | C8H23O4Si2 | Yes |
| Liver OAA332 | Oac | OAA332 | 332 | 1 2 3 4 | 4 | M+4 | C14H38NO5Si3 | No |
| Liver Akg346 | Akg.s | Akg346 | 346 | 1 2 3 4 5 | 5 | M+5 | C9H28O5NSi2 | Yes |
| Liver Malate391 | Mal.s | Malate391 | 391 | 2 3 4 | 3 | M+3 | C14H39O4Si3 | Yes |
| Liver Malate419 | Mal.s | Malate419 | 419 | 1 2 3 4 | 4 | M+4 | C14H39O5Si3 | Yes |
| Liver Asp302 | Asp.s | Asp302 | 302 | 1 2 | 2 | M+2 | C12H32O2NSi2 | Yes |
| Liver Asp390 | Asp.s | Asp390 | 390 | 2 3 4 | 3 | M+3 | C14H40O3NSi3 | Yes |
| Liver Asp418 | Asp.s | Asp418 | 418 | 1 2 3 4 | 4 | M+4 | C14H40O4NSi3 | Yes |
| Liver PEP453 | PEP | PEP453 | 453 | 1 2 3 | 3 | M+3 | C14H38O6Si3P | No |
| Liver DHAP484 | DHAP.s (Ignored) | DHAP484 | 484 | 1 2 3 | 3 | M+3 | C15H43O6NSi3P | No |
| Liver G3P571 | G3P | G3P571 | 571 | 1 2 3 | 3 | M+3 | C20H56O6Si4P | No |
| Liver Cit431 | Cit.s | Cit431 | 431 | 1 2 3 4 5 | 5 | M+5 | C14H39O5Si3 | No |
| Liver Cit459 | Cit.s | Cit459 | 459 | 1 2 3 4 5 6 | 6 | M+6 | C14H39O6Si3 | Yes |
| Perfusate Glc (DIO301) | Glc.perf | DIO301 | 301 | 1 2 3 4 5 6 | 6 | M+6 | C8H21O7 | No |
| Perfusate Glc (MOX145) | Glc.perf | MOX145 | 145 | 1 2 | 2 | M+2 | C4H11O3N | No |
| Perfusate Glc (Aldo173) | Glc.perf | Aldo173 | 173 | 5 6 | 2 | M+2 | C6H13O4 | No |
| Perfusate Glc (Aldo259) | Glc.perf | Aldo259 | 259 | 4 5 6 | 3 | M+3 | C9H19O6 | No |
| Perfusate Glc (Aldo284) | Glc.perf | Aldo284 | 284 | 1 2 3 4 | 4 | M+4 | C9H18O6N | No |
| Perfusate Glc (Aldo370) | Glc.perf | Aldo370 | 370 | 1 2 3 4 5 | 5 | M+5 | C12H24O8N | No |
| Effluent Glc (DIO301) | Glc.eff | DIO301 | 301 | 1 2 3 4 5 6 | 6 | M+6 | C8H21O7 | Yes |

| INCA measurement ID | Pool ID | Derivative / fragment | m/z | Labeled Atom IDs | Max labeled C | Max isotopologue | Unlabeled atoms | Active |
|---|---|---|---|---|---|---|---|---|
| Effluent Glc (MOX145) | Glc.eff | MOX145 | 145 | 1 2 | 2 | M+2 | C4H11O3N | Yes |
| Effluent Glc (Aldo173) | Glc.eff | Aldo173 | 173 | 5 6 | 2 | M+2 | C6H13O4 | Yes |
| Effluent Glc (Aldo259) | Glc.eff | Aldo259 | 259 | 4 5 6 | 3 | M+3 | C9H19O6 | Yes |
| Effluent Glc (Aldo284) | Glc.eff | Aldo284 | 284 | 1 2 3 4 | 4 | M+4 | C9H18O6N | Yes |
| Effluent Glc (Aldo370) | Glc.eff | Aldo370 | 370 | 1 2 3 4 5 | 5 | M+5 | C12H24O8N | Yes |
| Perfusate Pyr174 | Pyr.perf | Pyr174 | 174 | 1 2 3 | 3 | M+3 | C3H12O3NSi | No |
| Perfusate Lac233 | Lac.perf | Lac233 | 233 | 2 3 | 2 | M+2 | C8H25O2Si2 | Yes |
| Perfusate Lac261 | Lac.perf | Lac261 | 261 | 1 2 3 | 3 | M+3 | C8H25O3Si2 | Yes |
| Effluent Pyr174 | Pyr.eff | Pyr174 | 174 | 1 2 3 | 3 | M+3 | C3H12O3NSi | Yes |
| Effluent Lac233 | Lac.eff | Lac233 | 233 | 2 3 | 2 | M+2 | C8H25O2Si2 | Yes |
| Effluent Lac261 | Lac.eff | Lac261 | 261 | 1 2 3 | 3 | M+3 | C8H25O3Si2 | Yes |
| Liver Glu330 | Glu.s | Glu330 | 330 | 2 3 4 5 | 4 | M+4 | C12H36O2NSi2 | Yes |
| Liver Glu432 | Glu.s | Glu432 | 432 | 1 2 3 4 5 | 5 | M+5 | C14H42O4NSi3 | Yes |

#### S2F. Boundary-flux measurement errors

| Reaction | Standard deviation |
|---|---|
| 1 Total Glc IN | 0.0600 |
| 1 Total Glc OUT | 0.0250 |
| 1 Total Lac OUT | 2.4000 |
| 1 Total Pyr IN | 0.2400 |
| 1 Total Pyr OUT | 0.2400 |
| 1 Glycerol source | 0.1000 |
| Glycerol.eff | 0.1000 |
| 1 NA Lac IN | 1 |
| 1 Total Lac IN | 0.5000 |

### Table S3. Goodness of fit

*Lower and upper χ² bounds are for 95% confidence at the stated degrees of freedom (DOF). A fit was rejected only when the SSR exceeded the upper bound. * Excluded from flux analyses (see Methods). † Fitted without intracellular lactate (see Methods).*

#### S3A. Glucose fragment model, 0 mM glucose

*Datasets H0, J0 and K0 lacked the DIO m/z 301 fragment, reducing their DOF.*

| Dataset | State | SSR | Lower χ² | Upper χ² | DOF | Fit |
|---|---|---|---|---|---|---|
| A0 | Fasted | 14.5 | 7.6 | 30.2 | 17 | Within range |
| B0 | Fasted | 29.5 | 6.9 | 28.8 | 16 | Above upper bound |
| C0 | Fasted | 8.9 | 6.9 | 28.8 | 16 | Within range |
| D0 | Fasted | 32 | 7.6 | 30.2 | 17 | Above upper bound |
| F0 | Fasted | 8.9 | 7.6 | 30.2 | 17 | Within range |
| G0 | Fasted | 11.5 | 8.9 | 32.9 | 19 | Within range |
| H0 | Fasted | 287.8 | 3.8 | 21.9 | 11 | Above upper bound |
| I0 | Fasted | 64.1 | 6.9 | 28.8 | 16 | Above upper bound |
| J0 | Fasted | 28.4 | 2.7 | 19 | 9 | Above upper bound |
| K0 | Fasted | 53.8 | 0.8 | 12.8 | 5 | Above upper bound |
| L0 | Fasted | 30.6 | 3.8 | 21.9 | 11 | Above upper bound |
| M0 | Fed | 29.8 | 2.2 | 17.5 | 8 | Above upper bound |
| N0 | Fed | 48.3 | 2.7 | 19 | 9 | Above upper bound |
| O0 | Fed | 8.4 | 1.2 | 14.4 | 6 | Within range |
| P0 | Fed | 35.7 | 6.9 | 28.8 | 16 | Above upper bound |
| Q0 | Fed | 9.5 | 0.8 | 12.8 | 5 | Within range |
| R0 | Fed | 9.8 | 1.2 | 14.4 | 6 | Within range |
| S0 | Fed | 53.6 | 0.2 | 9.3 | 3 | Above upper bound |
| T0 | Fed | 1905.3 | 6.9 | 28.8 | 16 | Above upper bound |
| U0 | Fed | 360.3 | 4.4 | 23.3 | 12 | Above upper bound |
| V0 | Fed | 18.5 | 9.6 | 34.2 | 20 | Within range |
| W0 | Fed | 643.1 | 2.7 | 19 | 9 | Above upper bound |
| X0 | Fed | 119.3 | 1.7 | 16 | 7 | Above upper bound |

#### S3B. One-compartment model, full constraint set, 0 mM glucose

| Dataset | State | SSR | Lower χ² | Upper χ² | DOF | Fit |
|---|---|---|---|---|---|---|
| A0 | Fasted | 610.3 | 78.5 | 135.2 | 105 | Above upper bound |
| B0 | Fasted | 550.4 | 77.7 | 134.1 | 104 | Above upper bound |
| C0 | Fasted | 548.6 | 80.3 | 137.5 | 107 | Above upper bound |
| D0 | Fasted | 729.2 | 81.1 | 138.7 | 108 | Above upper bound |
| F0 | Fasted | 328.9 | 81.1 | 138.7 | 108 | Above upper bound |
| G0 | Fasted | 1022.6 | 82.9 | 140.9 | 110 | Above upper bound |
| H0 | Fasted | 933.9 | 75.9 | 131.8 | 102 | Above upper bound |
| I0 | Fasted | 855.1 | 80.3 | 137.5 | 107 | Above upper bound |
| J0 | Fasted | 525.8 | 74.2 | 129.6 | 100 | Above upper bound |
| K0 | Fasted | 427.7 | 68.2 | 121.6 | 93 | Above upper bound |
| L0 | Fasted | 233 | 75.9 | 131.8 | 102 | Above upper bound |
| M0 | Fed | 109.6 | 73.4 | 128.4 | 99 | Within range |
| N0 | Fed | 197.8 | 74.2 | 129.6 | 100 | Above upper bound |
| O0 | Fed | 94 | 71.6 | 126.1 | 97 | Within range |
| P0 | Fed | 114.7 | 80.3 | 137.5 | 107 | Within range |
| Q0 | Fed | 54 | 67.4 | 120.4 | 92 | Below lower bound |
| R0 | Fed | 149 | 69.1 | 122.7 | 94 | Above upper bound |
| S0 | Fed | 181.5 | 66.5 | 119.3 | 91 | Above upper bound |

| Dataset | State | SSR | Lower χ² | Upper χ² | DOF | Fit |
|---|---|---|---|---|---|---|
| T0 | Fed | 1960.4 | 78.5 | 135.2 | 105 | Above upper bound |
| U0 | Fed | 473.7 | 75.1 | 130.7 | 101 | Above upper bound |
| V0 | Fed | 127.8 | 83.7 | 142 | 111 | Within range |
| W0 | Fed | 430.3 | 72.5 | 127.3 | 98 | Above upper bound |
| X0 | Fed | 262.9 | 71.6 | 126.1 | 97 | Above upper bound |

#### S3C. One-compartment model, full constraint set, 5 mM glucose

| Dataset | State | SSR | Lower χ² | Upper χ² | DOF | Fit |
|---|---|---|---|---|---|---|
| A5 | Fasted | 684.9 | 168.1 | 247.6 | 206 | Above upper bound |
| B5 | Fasted | 554.2 | 167.2 | 246.5 | 205 | Above upper bound |
| C5 | Fasted | 884.7 | 163.6 | 242.2 | 201 | Above upper bound |
| D5 | Fasted | 1022 | 170 | 249.8 | 208 | Above upper bound |
| F5 | Fasted | 536.9 | 165.4 | 244.4 | 203 | Above upper bound |
| I5 | Fasted | 1715.8 | 161.8 | 240 | 199 | Above upper bound |
| J5 | Fasted | 583.7 | 167.2 | 246.5 | 205 | Above upper bound |
| M5 | Fed | 1893.4 | 195.3 | 280.4 | 236 | Above upper bound |
| N5 | Fed | 1269.8 | 180.8 | 263 | 220 | Above upper bound |
| Q5 | Fed | 917.3 | 160.9 | 238.9 | 198 | Above upper bound |
| R5 | Fed | 974.3 | 175.4 | 256.4 | 214 | Above upper bound |
| U5 | Fed | 1070.8 | 179.9 | 261.9 | 219 | Above upper bound |
| V5 | Fed | 1168.7 | 180.8 | 263 | 220 | Above upper bound |

#### S3D. One-compartment model, full constraint set, 20 mM glucose

| Dataset | State | SSR | Lower χ² | Upper χ² | DOF | Fit |
|---|---|---|---|---|---|---|
| G20* | Fasted | 31050.5 | 173.6 | 254.2 | 212 | Above upper bound |
| H20 | Fasted | 981.5 | 180.8 | 263 | 220 | Above upper bound |
| K20 | Fasted | 2512.2 | 170 | 249.8 | 208 | Above upper bound |
| L20 | Fasted | 2584.5 | 167.2 | 246.5 | 205 | Above upper bound |
| O20 | Fed | 897.8 | 164.5 | 243.3 | 202 | Above upper bound |
| P20 | Fed | 2802.6 | 168.1 | 247.6 | 206 | Above upper bound |
| S20 | Fed | 5162.6 | 166.3 | 245.4 | 204 | Above upper bound |
| T20 | Fed | 1874.4 | 167.2 | 246.5 | 205 | Above upper bound |
| W20 | Fed | 13572.1 | 181.7 | 264.1 | 221 | Above upper bound |
| X20 | Fed | 2428.4 | 167.2 | 246.5 | 205 | Above upper bound |

#### S3E. One-compartment model, reduced constraint set, 0 mM glucose

| Dataset | State | SSR | Lower χ² | Upper χ² | DOF | Fit |
|---|---|---|---|---|---|---|
| A0 | Fasted | 91.9 | 49.6 | 96.2 | 71 | Within range |
| B0 | Fasted | 59 | 48.8 | 95 | 70 | Within range |
| C0 | Fasted | 65.2 | 48.8 | 95 | 70 | Within range |
| D0 | Fasted | 94.8 | 49.6 | 96.2 | 71 | Within range |
| F0 | Fasted | 46.7 | 49.6 | 96.2 | 71 | Below lower bound |
| G0 | Fasted | 41 | 49.6 | 96.2 | 71 | Below lower bound |
| H0 | Fasted | 86.4 | 44.6 | 89.2 | 65 | Within range |
| I0 | Fasted | 77.7 | 48.8 | 95 | 70 | Within range |
| J0 | Fasted | 63.3 | 43 | 86.8 | 63 | Within range |
| K0 | Fasted | 85.6 | 37.2 | 78.6 | 56 | Above upper bound |
| L0 | Fasted | 83.5 | 44.6 | 89.2 | 65 | Within range |
| M0 | Fed | 42.7 | 42.1 | 85.7 | 62 | Within range |

| Dataset | State | SSR | Lower χ² | Upper χ² | DOF | Fit |
|---|---|---|---|---|---|---|
| N0 | Fed | 56.8 | 43 | 86.8 | 63 | Within range |
| O0 | Fed | 17.3 | 40.5 | 83.3 | 60 | Below lower bound |
| P0 | Fed | 51.8 | 48.8 | 95 | 70 | Within range |
| Q0 | Fed | 105 | 38.8 | 80.9 | 58 | Above upper bound |
| R0 | Fed | 15.8 | 40.5 | 83.3 | 60 | Below lower bound |
| S0 | Fed | 69.7 | 37.2 | 78.6 | 56 | Within range |
| T0 | Fed | 1930.5 | 48.8 | 95 | 70 | Above upper bound |
| U0 | Fed | 377 | 45.4 | 90.3 | 66 | Above upper bound |
| V0 | Fed | 91.7 | 52.1 | 99.7 | 74 | Within range |
| W0 | Fed | 161 | 41.3 | 84.5 | 61 | Above upper bound |
| X0 | Fed | 116.5 | 40.5 | 83.3 | 60 | Above upper bound |

#### S3F. One-compartment model, reduced constraint set, 5 mM glucose

| Dataset | State | SSR | Lower χ² | Upper χ² | DOF | Fit |
|---|---|---|---|---|---|---|
| A5 | Fasted | 206.2 | 99.5 | 162.3 | 129 | Above upper bound |
| B5 | Fasted | 176.3 | 97.7 | 160.1 | 127 | Above upper bound |
| C5 | Fasted | 93.8 | 93.3 | 154.5 | 122 | Within range |
| D5 | Fasted | 99.6 | 89.8 | 150 | 118 | Within range |
| F5 | Fasted | 127.5 | 94.2 | 155.6 | 123 | Within range |
| I5 | Fasted | 626.3 | 95.9 | 157.8 | 125 | Above upper bound |
| J5 | Fasted | 246.1 | 95.9 | 157.8 | 125 | Above upper bound |
| J5† | Fasted | 109 | 86.3 | 145.4 | 114 | Within range |
| M5 | Fed | 1586.2 | 95.1 | 156.7 | 124 | Above upper bound |
| N5 | Fed | 984.2 | 97.7 | 160.1 | 127 | Above upper bound |
| Q5 | Fed | 527.6 | 90.7 | 151.1 | 119 | Above upper bound |
| R5 | Fed | 600.3 | 94.2 | 155.6 | 123 | Above upper bound |
| U5 | Fed | 590.3 | 95.9 | 157.8 | 125 | Above upper bound |
| V5 | Fed | 573.5 | 97.7 | 160.1 | 127 | Above upper bound |

#### S3G. One-compartment model, reduced constraint set, 20 mM glucose

| Dataset | State | SSR | Lower χ² | Upper χ² | DOF | Fit |
|---|---|---|---|---|---|---|
| G20* | Fasted | 20075.6 | 103.8 | 167.9 | 134 | Above upper bound |
| H20 | Fasted | 194.7 | 109.1 | 174.6 | 140 | Above upper bound |
| K20 | Fasted | 1485.6 | 99.5 | 162.3 | 129 | Above upper bound |
| L20 | Fasted | 1969.3 | 97.7 | 160.1 | 127 | Above upper bound |
| O20 | Fed | 491.3 | 94.2 | 155.6 | 123 | Above upper bound |
| P20 | Fed | 1925.1 | 96.8 | 159 | 126 | Above upper bound |
| S20 | Fed | 3633.1 | 95.1 | 156.7 | 124 | Above upper bound |
| T20 | Fed | 1369.8 | 95.9 | 157.8 | 125 | Above upper bound |
| W20 | Fed | 10096.8 | 95.1 | 156.7 | 124 | Above upper bound |
| X20 | Fed | 1798.7 | 95.9 | 157.8 | 125 | Above upper bound |

#### S3H. Two-compartment model, 0 mM glucose

| Dataset | State | SSR | Lower χ² | Upper χ² | DOF | Fit |
|---|---|---|---|---|---|---|
| A0 | Fasted | 56.1 | 58 | 107.8 | 81 | Below lower bound |
| B0 | Fasted | 30.5 | 56.3 | 105.5 | 79 | Below lower bound |
| C0 | Fasted | 38 | 64.8 | 117 | 89 | Below lower bound |
| D0 | Fasted | 81.3 | 60.5 | 111.2 | 84 | Within range |
| F0 | Fasted | 16.6 | 60.5 | 111.2 | 84 | Below lower bound |

| Dataset | State | SSR | Lower χ² | Upper χ² | DOF | Fit |
|---|---|---|---|---|---|---|
| G0 | Fasted | 105.7 | 62.2 | 113.5 | 86 | Within range |
| H0 | Fasted | 118.6 | 55.5 | 104.3 | 78 | Above upper bound |
| I0 | Fasted | 116 | 58.8 | 108.9 | 82 | Above upper bound |
| J0 | Fasted | 85.1 | 52.9 | 100.8 | 75 | Within range |
| K0 | Fasted | 58.6 | 47.1 | 92.7 | 68 | Within range |
| L0 | Fasted | 44.3 | 55.5 | 104.3 | 78 | Below lower bound |
| M0 | Fed | 41.9 | 52.9 | 100.8 | 75 | Below lower bound |
| N0 | Fed | 71.3 | 53.8 | 102 | 76 | Within range |
| O0 | Fed | 26.8 | 51.3 | 98.5 | 73 | Below lower bound |
| P0 | Fed | 46.3 | 59.7 | 110.1 | 83 | Below lower bound |
| Q0 | Fed | 8.7 | 47.1 | 92.7 | 68 | Below lower bound |
| R0 | Fed | 33.2 | 47.9 | 93.9 | 69 | Below lower bound |
| S0 | Fed | 60.2 | 46.3 | 91.5 | 67 | Within range |
| T0 | Fed | 24.9 | 54.6 | 103.2 | 77 | Below lower bound |
| U0 | Fed | 7.5 | 40.5 | 83.3 | 60 | Below lower bound |
| V0 | Fed | 24.7 | 58 | 107.8 | 81 | Below lower bound |
| W0 | Fed | 144 | 52.1 | 99.7 | 74 | Above upper bound |
| X0 | Fed | 109.8 | 51.3 | 98.5 | 73 | Above upper bound |

#### S3I. Two-compartment model, 5 mM glucose

| Dataset | State | SSR | Lower χ² | Upper χ² | DOF | Fit |
|---|---|---|---|---|---|---|
| A5 | Fasted | 83.7 | 146.5 | 221.3 | 182 | Below lower bound |
| B5 | Fasted | 80.5 | 146.5 | 221.3 | 182 | Below lower bound |
| C5 | Fasted | 107.6 | 132.2 | 203.6 | 166 | Below lower bound |
| D5 | Fasted | 90.7 | 138.5 | 211.3 | 173 | Below lower bound |
| F5 | Fasted | 86.5 | 143.8 | 217.9 | 179 | Below lower bound |
| I5 | Fasted | 112.2 | 140.3 | 213.5 | 175 | Below lower bound |
| J5 | Fasted | 87.3 | 145.6 | 220.1 | 181 | Below lower bound |
| M5 | Fed | 111.1 | 174.5 | 255.3 | 213 | Below lower bound |
| N5 | Fed | 53.2 | 160 | 237.8 | 197 | Below lower bound |
| Q5 | Fed | 65.1 | 140.3 | 213.5 | 175 | Below lower bound |
| R5 | Fed | 70.9 | 154.6 | 231.2 | 191 | Below lower bound |
| U5 | Fed | 97.8 | 159.1 | 236.7 | 196 | Below lower bound |
| V5 | Fed | 97.6 | 160 | 237.8 | 197 | Below lower bound |

#### S3J. Two-compartment model, 20 mM glucose

| Dataset | State | SSR | Lower χ² | Upper χ² | DOF | Fit |
|---|---|---|---|---|---|---|
| G20* | Fasted | 1556.3 | 157.3 | 234.5 | 194 | Above upper bound |
| H20 | Fasted | 102.6 | 159.1 | 236.7 | 196 | Below lower bound |
| K20 | Fasted | 152.8 | 148.3 | 223.5 | 184 | Within range |
| L20 | Fasted | 117.4 | 145.6 | 220.1 | 181 | Below lower bound |
| O20 | Fed | 37.2 | 143.8 | 217.9 | 179 | Below lower bound |
| P20 | Fed | 57.1 | 147.4 | 222.4 | 183 | Below lower bound |
| S20 | Fed | 88.3 | 145.6 | 220.1 | 181 | Below lower bound |
| T20 | Fed | 56.4 | 146.5 | 221.3 | 182 | Below lower bound |
| W20 | Fed | 139.7 | 160.9 | 238.9 | 198 | Below lower bound |
| X20 | Fed | 96.2 | 146.5 | 221.3 | 182 | Below lower bound |
